# Peatland *Acidobacteria* with a dissimilatory sulfur metabolism

**DOI:** 10.1101/197269

**Authors:** Bela Hausmann, Claus Pelikan, Craig W. Herbold, Stephan Köstlbacher, Mads Albertsen, Stephanie A. Eichorst, Tijana Glavina del Rio, Martin Huemer, Per H. Nielsen, Thomas Rattei, Ulrich Stingl, Susannah G. Tringe, Daniela Trojan, Cecilia Wentrup, Dagmar Woebken, Michael Pester, Alexander Loy

## Abstract

Sulfur-cycling microorganisms impact organic matter decomposition in wetlands and consequently greenhouse gas emissions from these globally relevant environments. However,their identities and physiological properties are largely unknown. By applying a functional metagenomics approach to an acidic peatland, we recovered draft genomes of seven novel *Acidobacteria* species with the potential for dissimilatory sulfite (*dsrAB*, *dsrC*, *dsrD*, *dsrN, dsrT, dsrMKJOP*) or sulfate respiration (*sat, aprBA, qmoABC* plus *dsr* genes). Surprisingly, the genomes also encoded *dsrL*, a unique gene of the sulfur oxidation pathway. Metatranscriptome analysis demonstrated expression of acidobacterial sulfur-metabolism genes in native peat soil and their upregulation in diverse anoxic microcosms. This indicated an active sulfate respiration pathway, which, however, could also operate in reverse for sulfur oxidation as recently shown for other microorganisms. *Acidobacteria* that only harbored genes for sulfite reduction additionally encoded enzymes that liberate sulfite from organosulfonates, which suggested organic sulfur compounds as complementary energy sources. Further metabolic potentials included polysaccharide hydrolysis and sugar utilization, aerobic respiration, several fermentative capabilities, and hydrogen oxidation. Our findings extend both, the known physiological and genetic properties of *Acidobacteria* and the known taxonomic diversity of microorganisms with a DsrAB-based sulfur metabolism, and highlight new fundamental niches for facultative anaerobic *Acidobacteria* in wetlands based on exploitation of inorganic and organic sulfur molecules for energy conservation.

## Introduction

Specialized microorganisms oxidize, reduce, or disproportionate sulfur compounds of various oxidation states (– II to + VI) to generate energy for cellular activity and growth and thereby drive the global sulfur cycle. The capability for characteristic sulfur redox reactions such as dissimilatory sulfate reduction or sulfide oxidation is not confined to single taxa but distributed across different, often unrelated taxa. The true extent of the taxon-diversity within the different guilds of sulfur microorganisms is unknown (Wasmund *et al.*, 2017). However, ecological studies employing specific sulfur metabolism genes (e.g., dissimilatory adenylyl-sulfate reductaseencoding *aprBA*, dissimilatory sulfite reductase-encoding *dsrAB*, or *soxB* that codes for a part of the thiosulfate-oxidizing Sox enzyme machinery) as phylogenetic and functional markers have repeatedly demonstrated that only a minor fraction of the sulfur metabolism gene diversity in many environments can be linked to recognized taxa (Meyer *et al.*, 2007; Müller *et al.*, 2015; Watanabe *et al.*, 2016). A systematic review of *dsrAB* diversity has revealed that the reductive bacterial-type enzyme branch of the DsrAB tree contains at least thirteen family-level lineages without any cultivated representatives. This indicates that major taxa of sulfate-/ sulfite-reducing microorganisms have not yet been identified (Müller *et al.*, 2015).

Wetlands are among those ecosystems that harbor a diverse community of microorganisms with reductive-type DsrAB, most of which cannot be identified because they are distant from recognized taxa (Pester *et al.*, 2012). Sulfur-cycling microorganisms provide significant ecosystem services in natural and anthropogenic wetlands, which are major sources of the climate-warming greenhouse gas methane (Kirschke *et al.*, 2013; Saunois *et al.*, 2016). While inorganic sulfur compounds are often detected only at low concentration (lower μM range), fast sulfur cycling nevertheless ensures that oxidized sulfur compounds such as sulfate are rapidly replenished for anaerobic respiration. The activity of sulfate-reducing microorganisms (SRM) fluctuates with time and space, but at peak times can account for up to 50% of anaerobic mineralization of organic carbon in wetlands (Pester *et al.*, 2012). Simultaneously, SRM prevent methane production by rerouting carbon flow away from methanogenic archaea. Autochthonous peat microorganisms that are affiliated to known SRM taxa, such as *Desulfosporosinus, Desulfomonile*, and *Syntrophobacter*, are typically low abundant (Loy *et al.*, 2004; Costello and Schmidt, 2006; Dedysh *et al.*, 2006; Kraigher *et al.*, 2006; Pester *et al.*, 2010; Steger *et al.*, 2011; Tveit *et al.*, 2013; Hausmann *et al.*, 2016). In contrast, some microorganisms that belong to novel, environmental *dsrAB* lineages can be considerably more abundant in wetlands than species-level *dsrAB* operational taxonomic units of known taxa (Steger *et al.*, 2011). However, the taxonomic identity of these novel *dsrAB*-containing microorganisms and their role in sulfur and carbon cycling has yet to be revealed.

Here, we recovered thirteen metagenome-assembled genomes (MAGs) encoding DsrAB of uncultured species from a peat soil by using a targeted, functional metagenomics approach. We analyzed expression of predicted physiological capabilities of the MAGs by metatranscriptome analyses of anoxic peat soil microcosms that were periodically stimulated by small additions of individual fermentation products with or without supplemented sulfate (Hausmann *et al.*, 2016). Our results suggest that some facultatively anaerobic members of the diverse *Acidobacteria* community in wetlands employ a dissimilatory sulfur metabolism.

## Materials and methods

### Anoxic microcosm experiments, stable isotope probing, and nucleic acids isolation

DNA and RNA samples were retrieved from a previous peat soil microcosm experiment (Hausmann *et al.*, 2016). Briefly, soil from 10 – 20 cm depth was sampled from an acidic peatland (Schlöppnerbrunnen II, Germany) in September 2010, and stored at 4 °C for one week prior to nucleic acids extractions and set-up of soil slurry incubations. Individual soil slurry microcosms were incubated anoxically (100% N_2_ atmosphere) in the dark at 14 °C, and regularly amended with low amounts (<200 μM) of either formate, acetate, propionate, lactate, butyrate, or without any additional carbon sources (six replicates each). In addition, half of the microcosms for each substrate were periodically supplemented with low amounts of sulfate (initial amendment of 190 – 387 μM with periodic additions of 79 – 161 μM final concentrations). DNA and RNA were extracted from the native soil and RNA was additionally extracted from the soil slurries after 8 and 36 days of incubation.

Furthermore, DNA was obtained from the heavy, ^13^C-enriched DNA fractions of a previous DNAstable isotope probing (DNA-SIP) experiment with soil from the same site (Pester *et al.*, 2010). Analogous to the single-substrate incubations, anoxic soil slurries were incubated for two months with low-amounts of sulfate and a ^13^C-labelled mixture of formate, acetate, propionate, and lactate. DNA was extracted, separated on eight replicated density gradients, and DNA from a total of 16 heavy fractions (density 1.715 – 1.726 g mL^-1^) was pooled for sequencing.

Additional DNA was obtained from soils that were sampled from different depths in the years 2004 and 2007 (Steger *et al.*, 2011).

### Quantitative PCR and metagenome/-transcriptome sequencing

Abundances of *Acidobacteria* subdivision 1, 2, and 3 in soil samples from different years and depths were determined by newly-developed 16S rRNA gene-targeted real-time quantitative PCR (qPCR) assays (Supplementary Methods). Native soil DNA (two libraries), heavy 13Cenriched DNA (three libraries), and native soil RNA, and RNA samples from the microcosms were sequenced on an Illumina HiSeq2000 system (Supplementary Methods).

### Binning, phylogeny, and annotation of DsrAB-encoding genomes

The differential coverage binning approach by Albertsen *et al.* (2013) was applied to extract MAGs of interest. The raw FASTQ paired-end reads were imported into the CLC Genomics Workbench 5.5.1 (CLC Bio) and trimmed using a minimum Phred quality score of 20 with no ambiguous nucleotides allowed. TruSeq adapters were removed and a minimum length filter of 50 nt was applied. This resulted in 214, 171, 233, 49, and 294 million reads after quality filtering and trimming for the two native soil and three SIP metagenomes, respectively (84 – 95% of the raw reads). All reads were co-assembled using CLCs *de novo* assembly algorithm (kmer size 63, bubble size 50, minimum scaffold size 1000 nt). The reads from all five metagenomes were independently mapped to the assembled scaffolds using CLCs map to reference function (minimum length 0.7, minimum similarity 0.95) to obtained the scaffold coverage. The SIP metagenomes were merged into one mapping. 137, 112, and 376 million reads could be mapped to the two native soil metagenomes and the SIP metagenome, respectively (64 – 66% of quality filtered reads). Gene prediction of the complete assembly was performed using prodigal (Hyatt *et al.*, 2010). In addition to the detection and taxonomic classification of 105 essential marker genes (Albertsen *et al.*, 2013), *dsrA* and *dsrB* homologs were identified using TIGRFAM’s hidden Markov model (HMM) profiles TIGR02064 and TIGR02066, respectively, with HMMER 3.1 (Eddy, 2011) and the provided trusted cut-offs. Additional *dsrAB*-containing scaffolds were identified by using tblastn with the published DsrAB database as a query against the assembly (Müller *et al.*, 2015). DsrAB sequences were classified by phylogenetic analysis (Supplementary Methods; Müller *et al.*, 2015). Binning and decontamination was finalized utilizing the G + C content and tetramer frequencies of the scaffolds, as well as paired-end information, as described and recommended in Albertsen *et al.* (2013). Completeness, contamination, and strain heterogeneity was estimated using CheckM 1.0.6 (Parks *et al.*, 2015) with lineage-specific marker sets selected at phylum rank (or class rank for *Proteobacteria*). MAGs were taxonomically classified by phylogenomic analysis of concatenated marker sequences and calculation of average nucleic and amino acid identities (ANI, AAI, Supplementary Methods). MAGs were annotated using MaGe (Vallenet *et al.*, 2017) and eggNOG (Huerta-Cepas *et al.*, 2016). Genes of interest (Supplementary Table S2) were manually curated using the full range of tools integrated in MaGe (Supplementary Methods).

### Genome-centric activity analysis: iRep and metatranscriptomics

The index of replication (iRep) was calculated for each MAG with the combined native soil metagenomes. Settings and thresholds were applied as recommended (Brown *et al.*, 2016) using bowtie2 (Langmead and Salzberg, 2012) and the iRep script with default settings. Qualityfiltered metatranscriptome reads were mapped to all genomes using bowtie2 and counted with featureCounts (Liao *et al.*, 2014). To determine gene expression changes, we applied the DESeq2 pipeline with recommended settings (Love *et al.*, 2014) (Supplementary Methods).

### Data availability

Metagenomic and -transcriptomic data were deposited under the BioProject accession numbers PRJNA412436 and PRJNA412438, respectively, and can also be obtained via the JGI’s genome portal (JGI Proposal ID 605). MAGs were deposited under the BioProject accession number PRJNA412580.

## Results

### Functional metagenomics: Recovery of *dsrAB*-containing acidobacterial genomes from native soil and ^13^C-DNA fraction metagenomes

This study was conducted with soil samples from the Schlöppnerbrunnen II peatland in Germany, which is a long-term study site with active sulfur cycling and harbors a large diversity of unknown microorganisms with divergent *dsrAB* genes (Steger *et al.*, 2011; Pester *et al.*, 2012). We initially generated co-assembled metagenomes from native peat soil DNA (53 Gb) and a pool of DNA extracts from the heavy fractions of a previous DNA-stable isotope probing (DNA-SIP) experiment with soil from the same peat (101 Gb). The heavy fractions, which were obtained from anoxic peat incubations with periodically supplemented sulfate and a mixture of **^13^**C-labelled formate, acetate, propionate, and lactate at low concentrations, were enriched in DNA from *Desulfosporosinus* and also harbored DNA from yet unidentified *dsrAB*-containing microorganisms (Pester *et al.*, 2010). Based on the metagenome data, the native peat was dominated by *Acidobacteria* (61%), but also had *Actinobacteria, Alphaproteobacteria*, and *Deltaproteobacteria* as abundant (>5%) phyla / classes (Figure 1). Dominance of *Acidobacteria, Alpha -* and *Deltaproteobacteria* is typical for peatlands (Dedysh, 2011). Quantitative PCR confirmed that *Acidobacteria* subdivisions 1, 2, and 3 persistently dominated the Schlöppnerbrunnen II peat microbiota in oxic and anoxic soil layers (Supplementary Methods, Figure 1), as observed in other peatlands (Serkebaeva *et al.*, 2013; Urbanová and Bárta, 2014; Ivanova *et al.*, 2016).

**Figure 1.**
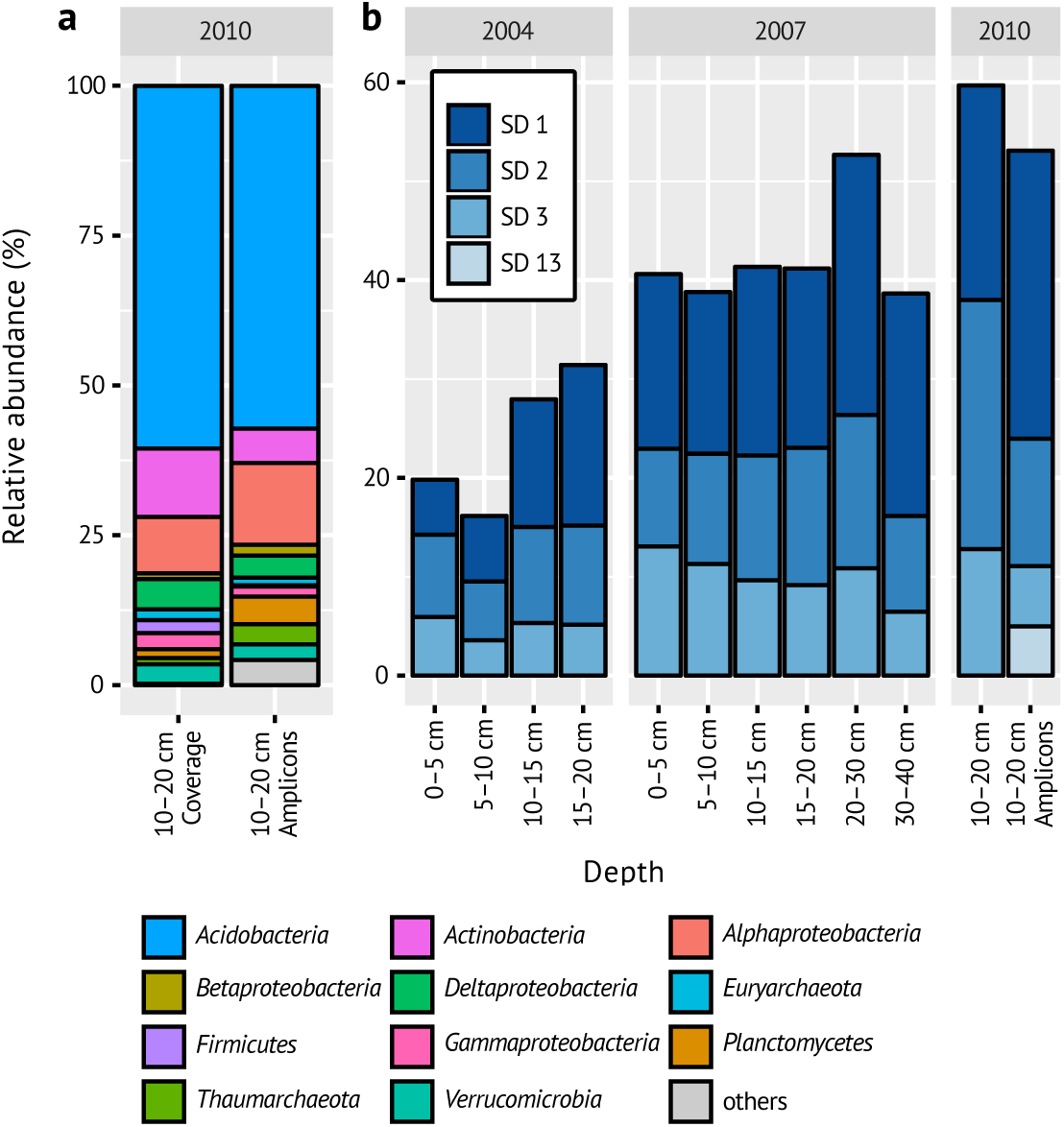
Microbial community composition in Schlöppnerbrunnen II peatland in samples from different years and soil depths. (a) Abundance of phyla and proteobacterial classes in the native soil (relative to all classified reads / amplicons). Taxa less abundant than 1% are grouped in grey. Coverage abundance is based on metagenomic reads mapped to classified scaffolds. Amplicon abundance is based on *rrn* operon-copy number-corrected abundance of 16S rRNA gene operational taxonomic units (Hausmann *et al.*, 2016). (b) Relative abundance of acidobacterial subdivisions (SD) in the native soil samples as determined by 16S rRNA gene qPCR assays. In addition, all subdivisions more abundant than 1% in a 16S rRNA gene amplicon dataset are shown (Hausmann *et al.*, 2016).

We identified 36 complete or partial *dsrAB* genes on scaffolds of the composite metagenome and subsequently recovered thirteen MAGs of DsrAB-encoding bacteria by differential coverage binning (Supplementary Table S1, Albertsen *et al.*, 2013). Twenty-eight *dsrAB* sequences were part of the reductive bacterial-type DsrAB family branch and were closely related to previously recovered sequences from this and other wetlands (Supplementary Figure S1). These *dsrAB* sequences were affiliated to the known SRM genera *Desulfosporosinus* (*Firmicutes*, n = 1, one MAG) and *Syntrophobacter* (*Deltaproteobacteria*, n = 3, two MAGs), the *Desulfobacca acetoxidans* lineage (n = 1), and the uncultured DsrAB family-level lineages 8 (n = 19, seven MAGs) and 10 (n = 4). Six sequences grouped with the oxidative bacterial-type DsrAB family and were distantly affiliated with *Sideroxydans lithotrophicus* (*Betaproteobacteria*, n = 5, two MAGs) or *Rhodomicrobium vannielii* (*Alphaproteobacteria*, n = 1) (Supplementary Figure S2). Interestingly, two of our sequences (n = 2, one MAG) and a DsrAB sequence from the candidate phylum *Rokubacteria* (Hug *et al.*, 2016) formed a completely novel basal lineage outside the four previously recognized DsrAB enzyme families (Supplementary Figure S2) (Müller *et al.*, 2015). The thirteen partial to near complete *dsrAB*-containing MAGs had moderate to no detectable contamination as assessed by CheckM (Supplementary Table S1) (Parks *et al.*, 2015) and derived from *Acidobacteria* subdivisions 1 and 3 (SbA1 – 7), *Desulfosporosinus* (SbF1), *Syntrophobacter* (SbD1, SbD2), *Betaproteobacteria* (SbB1, SbB2), and *Verrucomicrobia* (SbV1), as inferred by phylogenetic analysis of DsrAB sequences (Supplementary Figures S1 and S2) and concatenated sequences of single-copy, phylogenetic marker genes (Supplementary Figure S3). Only the *Desulfosporosinus* and *Syntrophobacter* MAGs contained rRNA gene sequences.

Phylogenomic analysis showed that *Acidobacteria* MAGs SbA1, SbA5, and SbA7 are affiliated with subdivision 1, while SbA3, SbA4, and SbA6 are affiliated with subdivision 3 (Supplementary Figure S3). The partial MAG SbA2 lacked the marker genes used for phylogenomic treeing, but was unambiguously assigned to *Acidobacteria* using an extended marker gene set (Albertsen *et al.*, 2013) and DsrAB phylogeny. The two near complete (96%) MAGs SbA1 and SbA5 have a size of 5.4 and 5.3 Mb, respectively. The G + C content of all MAGs ranges from 58% to 63% (Supplementary Table S1). This in accordance with genome characteristics of acidobacterial isolates, which have genome sizes of 4.1 – 10.0 Mb and G + C contents of 57 – 62% (Ward *et al.*, 2009; Rawat *et al.*, 2012,). SbA1 and SbA7 form a monophyletic clade in the *Acidobacteria* subdivision 1 with an AAI (Rodriguez-R and Konstantinidis, 2014) of 63% (Supplementary Figure S3) and DsrAB identity of 80%. They have 56% AAI to their closest relative, *Ca. Koribacter versatilis*, which is lower than AAIs among members of known acidobacterial genera (60 – 71%). The third MAG from subdivision 1, SbA5, is affiliated with *Terracidiphilus gabretensis* with an AAI of 61%. DsrAB identity of SbA5 to SbA1 and SbA7 is 79%. The three subdivision 3 MAGs form a monophyletic clade with *Ca. Solibacter usitatus* (Supplementary Figure S3). SbA3, SbA4, and SbA6 have AAIs of 59 – 73% amongst them and 61 – 62% to *Ca. S. usitatus*. DsrAB identity amongst the three MAGs is 80 – 94% and 74 – 79% to the subdivisions 1 MAGs.

The DsrAB sequences encoded on all seven MAGs are affiliated with the uncultured DsrAB family-level lineage 8 (Supplementary Figure S1), which so far only consisted of environmental *dsrAB* sequences of unknown taxonomic identity (Müller *et al.*, 2015). Based on these MAGs and metatranscriptome analyses of anoxic peat soil microcosms, we here describe the putative metabolic capabilities of these novel DsrAB-encoding *Acidobacteria*. Details on the other MAGs will be described elsewhere (Hausmann *et al.*, unpublished; Anantharaman *et al.*, unpublished). Functional interpretations of the recovered MAGs are made under the premise that the genomes are not closed, and thus it is unknown if genes are absent in these organisms or are missing due to incomplete sequencing, assembly, or binning.

### Dissimilatory sulfur metabolism

Although *Acidobacteria* are abundant in diverse environments with active sulfur cycling (Serkebaeva *et al.*, 2013; Urbanová and Bárta, 2014; Sánchez-Andrea *et al.*, 2011; Wang *et al.*, 2012), this is the first discovery of members of this phylum with a putative dissimilatory sulfur metabolism. SbA2, SbA3, and SbA7 encode the complete canonical pathway for dissimilatory sulfate reduction, including homologs for sulfate transport (*sulP* and / or dass, not in SbA7) and activation (*sat*, ppa, *hppA*), adenosine 5′-phosphosulfate (APS) reduction (*aprBA, qmoABC*), and sulfite reduction (*dsrAB, dsrC, dsrMKJOP*) (Figure 2, Supplementary Table S2a) (Santos *et al.*, 2015). SbA1, SbA4, SbA5, and SBA6 have an incomplete sulfate reduction gene set but contain all *dsr* genes for sulfite reduction. Several other *dsr* genes were present on some of the MAGs. The *dsrD* and *dsrN* genes occurred in pairs. The role of the small DsrD protein is unresolved, but its ubiquity among SRM suggests an essential function in sulfate reduction (Hittel and Voordouw, 2000). DsrN is a homolog of cobyrinate a,c-diamide synthase in cobalamin biosynthesis and may be involved in amidation of the siroheme prosthetic group of DsrAB (Lübbe *et al.*, 2006). DsrV, a homolog of precorrin-2 dehydrogenase, and DsrWa, a homolog of uroporphyrin-III C-methyltransferase, may also be involved in siroheme biosynthesis (Holkenbrink *et al.*, 2011). DsrT is required for sulfide oxidation in *Chlorobaculum tepidum*, but also found in SRM (Holkenbrink *et al.*, 2011). The presence of *dsrMK*-paralogs (*dsrM2, dsrK2*) upstream of *dsrAB* is not uncommon in SRM (Pereira *et al.*, 2011). DsrMK are present in all *dsrAB*-containing microorganisms and are a transmembrane module involved in reduction of cytoplasmic DsrC-trisulfide in SRM, the final step in sulfate reduction (Santos *et al.*, 2015). DsrC encoded on the MAGs have the two essential cysteine residues at the C-terminal end for full functionality (Venceslau *et al.*, 2014). Interestingly, *dsrC* forms a gene duo with *dsrL* downstream of *dsrAB* in all seven MAGs. This is surprising, because *dsrL* is not found in SRM but in sulfur oxidizers. DsrL is highly expressed and essential for sulfur oxidation by the purple sulfur bacterium *Allochromatium vinosum* (Lübbe *et al.*, 2006; Weissgerber *et al.*, 2014). DsrL is a cytoplasmic iron-sulfur flavoprotein with proposed NAD(P)H:acceptor oxidoreductase activity and was copurified with DsrAB from the soluble fraction of A. vinosum (Dahl *et al.*, 2005). Given the possible role of DsrL in sulfur oxidation, we sought to detect additional genes indicative of oxidative sulfur metabolism in the acidobacterial MAGs. However, genes for sulfide:quinone reductase (*sqr*), adenylyl-sulfate reductase membrane anchor subunit (*aprM*), flavocytochrome c sulfide dehydrogenase (*fccAB*), sulfur reductase (*sreABC*), thiosulfate reductase (*phsABC*), polysulfide reductase (*psrABC*), membrane-bound sulfite oxidizing enzyme (*soeABC*), cytoplasmic sulfur trafficking enzymes (*tusA, dsrE2, dsrEFH*), or DsrQ / DsrU (unknown functions) were absent (Laska *et al.*, 2003; Holkenbrink *et al.*, 2011; Lenk *et al.*, 2012; Wasmund *et al.*, 2017). SbA1, SbA3, SbA4, and SbA6 contain genes that have only low homology to *soxCD* / *sorAB*, periplasmic sulfite-oxidizing enzymes (Supplementary Results) and, thus, might have another function (Ghosh and Dam, 2009).

**Figure 2.**
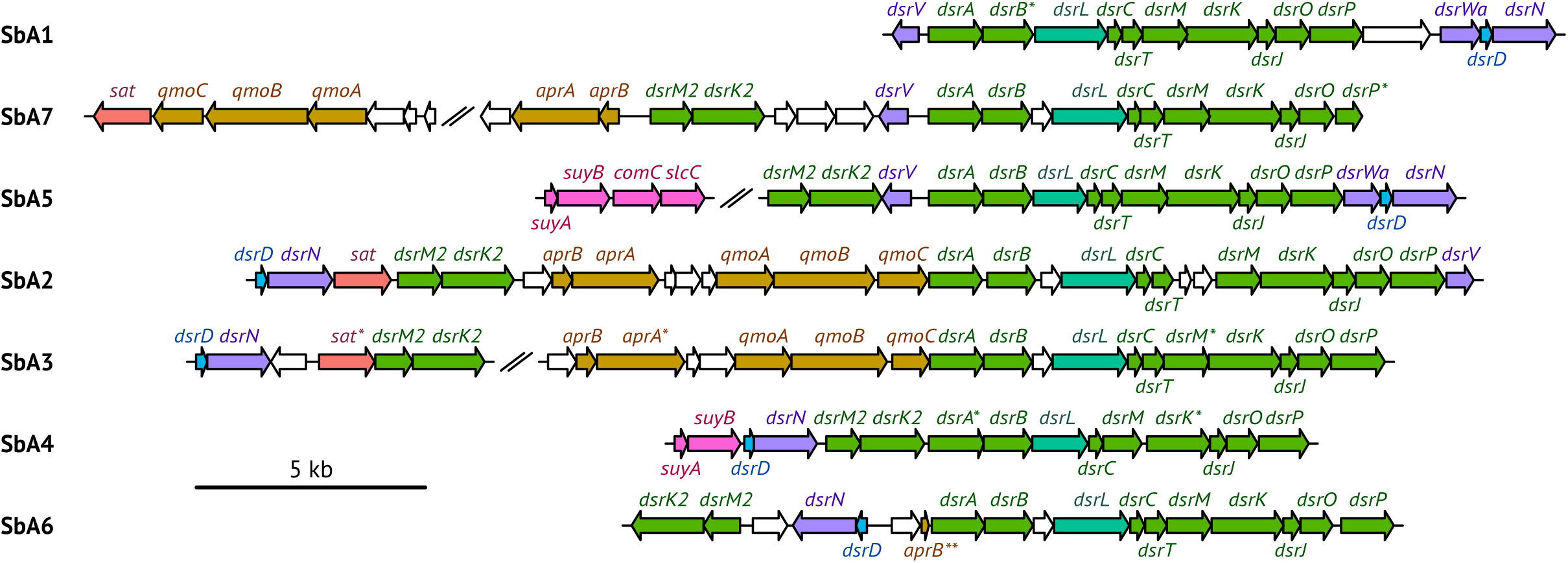
Organization of dissimilatory sulfur metabolism genes on acidobacterial MAGs SbA1 – 7. Red: *sat*; orange: *aprBA, qmoABC*; green: *dsrABCMKJOPM2K2*; blue: *dsrD*; turquoise: *dsrL*; violet: *dsrNVWa*; pink: *suyAB, comC, slcC*; white: genes of unknown function or not involved in sulfur metabolism. In SbA2 all genes are on one scaffold (scaffold 0lkb). Gene fragments at contig borders are indicated by an asterisk. *aprB* in SbA6, indicated by two asterisks, is truncated, which indicates a pseudogene or is due to an assembly error. Scaffolds are separated by two slashes.

Despite ongoing sulfur cycling, concentrations of inorganic sulfur compounds such as sulfate are low (lower μM range) in the Schlöppnerbrunnen II peatland (Schmalenberger *et al.*, 2007; Küsel *et al.*, 2008; Knorr and Blodau, 2009). Enzymatic release of inorganic sulfur compounds from organic matter might thus represent a significant resource for sulfur-dissimilating microorganisms. We thus specifically searched for genes coding for known organosulfur reactions that yield sulfite (Wasmund *et al.*, 2017). Genes for cysteate sulfo-lyase (*cuyA*), methanesulfonate monooxygenase (*msmABCD*), sulfoacetaldehyde acetyltransferase (xsc), and taurine dioxygenase (*tauD*) were absent. However, *suyAB*, coding for the (*R*)-sulfolactate sulfolyase complex that cleaves (*R*)-sulfolactate into pyruvate and sulfite (Denger and Cook, 2010), were present in SbA4 and SbA5 (Supplementary Table S2a). Intriguingly, SbA4 and SbA5 only have capability for sulfite reduction. SbA5 also encodes the racemase machinery for (*S*)-sulfolactate to (*R*)-sulfolactate, (*S*)-sulfolactate dehydrogenase (*slcC*) and (*R*)-sulfolactate dehydrogenase (*comC*); the regulator gene *suyR* or the putative importer SlcHFG were absent (Denger and Cook, 2010). Pyruvate may be used as an energy and carbon source, while sulfite could be used as an electron acceptor for anaerobic respiration (Simon and Kroneck, 2013).

### Respiration

Cultivated *Acidobacteria* of subdivisions 1 and 3 are strict aerobes or facultative anaerobes (e.g., Eichorst *et al.*, 2007; Kulichevskaya *et al.*, 2010, 2014; Pankratov and Dedysh, 2010; Dedysh *et al.*, 2012; Pankratov *et al.*, 2012). Accordingly, we found respiratory chains encoded in all acidobacterial MAGs (Figure 3, Supplementary Results), with (near) complete operons for NADH dehydrogenase 1, succinate dehydrogenase (lacking in SbA2), one or both types of quinol — cytochrome-c reductase, low-affinity terminal oxidases, and ATP synthase (lacking in SbA2) (Supplementary Tables S2b – h). High-affinity terminal oxidases, putatively involved in detoxification of oxygen (Ramel *et al.*, 2013; Giuffrè *et al.*, 2014), are limited to four MAGs (Supplementary Table S2g). Genes for dissimilatory nitrogen or iron metabolisms are absent, with the exception of a putative metal reductase in SbA2 of unclear physiological role (Supplementary Results).

**Figure 3.**
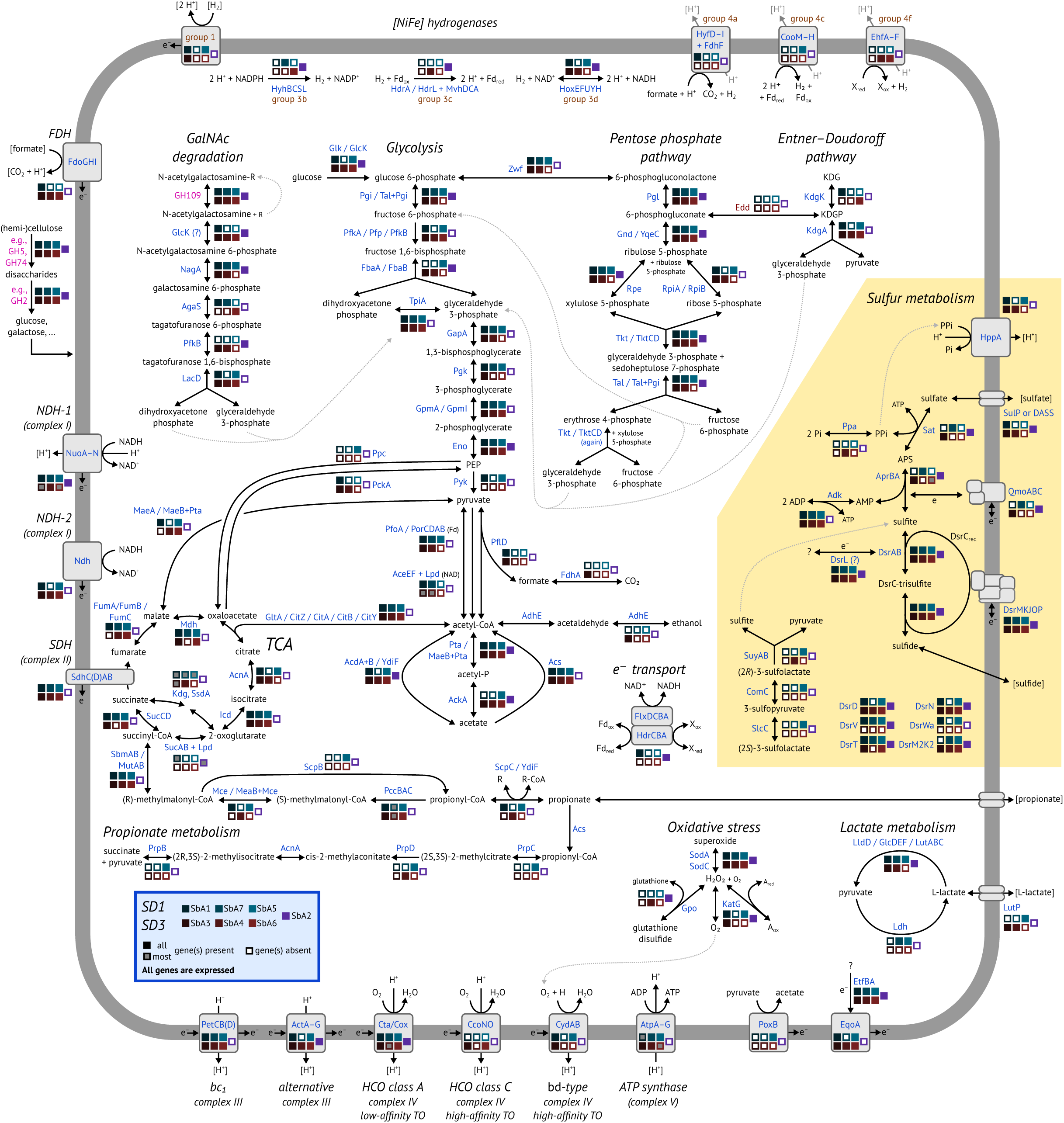
Metabolic model as inferred from analysis of acidobacterial MAGs SbA1 – 7. Sulfur metabolism is highlighted in yellow. Enzymes and transporters are shown in blue font. Glycoside hydrolases are shown in pink font (Supplementary Table S2). Extracellular compounds are in parentheses. A slash (/) indicates isozymes, i.e., enzymes that perform the same function, but are distinctly different or have more than one established name. AcdA + B, MaeB + Pta, MeaB + Mce, Tal + Pgi: bifunctional fusion genes / proteins. Otherwise the plus sign (+) indicates protein complexes. TCA: tricarboxylic acid cycle. FDH: formate dehydrogenase. Hase: hydrogenase. NDH: NADH dehydrogenase. HCO: haem-copper oxidase. TO: terminal oxidase. KDG: 2-dehydro-3-deoxy-Dgluconate. KDGP: 2-dehydro-3-deoxy-D-gluconate 6-phosphate. Expression of at least one copy of every enzyme and transporter was observed in the incubation samples.

### Hydrogen utilization and production

We identified [NiFe] hydrogenases of groups 1, 3, and 4 (Greening *et al.*, 2016) in SbA1 – 7 (Supplementary Table S2j). Membrane-bound group 1 hydrogenases (SbA1, SbA3, SbA5) consume hydrogen from the periplasm as an electron donor to generate energy, possibly coupled to sulfate / sulfite reduction. In contrast to other *Acidobacteria*, no group 1h / 5 hydrogenases, which are coupled to oxygen respiration, were identified (Greening *et al.*, 2015). Cytoplasmic group 3 hydrogenases (all MAGs) are bidirectional and proposed to be involved in energy-generating hydrogen oxidation and / or fermentative hydrogen production. Membranebound group 4 hydrogenases (SbA1, SbA5, SbA4, SbA6) produce H_2_ and are postulated to conserve energy by proton translocation by oxidizing substrates like formate (group 4a) or carbon monoxide (via ferredoxin, group 4c) (Figure 3).

### A versatile heterotrophic physiology

*Acidobacteria* are known for their capability to degrade simple and polymeric carbohydrates (e.g., Kulichevskaya *et al.*, 2010, 2014; Pankratov and Dedysh, 2010; Dedysh *et al.*, 2012; Eichorst *et al.*, 2011; Pankratov *et al.*, 2012; Rawat *et al.*, 2012; Huber *et al.*, 2016), supported by many diverse carbohydrate-active enzymes encoded in their genomes (Ward *et al.*, 2009; Rawat *et al.*, 2012). Accordingly, the MAGs recovered in our study also contain many genes encoding diverse carbohydrate-active enzymes (Supplementary Methods, Figure 4). These include glycoside hydrolases (GH, 1.0 – 4.0% of all genes), polysaccharide lyases (0.07 – 0.3%), and carbohydrate esterases (0.7 – 1.4%) that are generally involved in degradation of complex sugars, but also glycosyltransferases (0.9 – 1.4%) for biosynthesis of carbohydrates. Functional GH gene groups (assigned by EC number) involved in cellulose and hemicellulose degradation were most prevalent (Supplementary Table S4). Specifically, the most abundant EC numbers are involved in cellulose (EC 3.2.1.4, e.g., GH5, GH74), xyloglucan (EC 3.2.1.150, EC 3.2.1.151, e.g., GH5, GH74), or xylan (EC 3.2.1.8, EC 3.2.1.37, e.g., GH5) degradation, similar to other members of *Acidobacteria* subdivision 1 and 3 (Ward *et al.*, 2009; Rawat *et al.*, 2012). Other abundant EC numbers were associated with oligosaccharide degradation (EC 3.2.1.21, e.g., GH2) or α-*N*acetylgalactosaminidase genes (EC 3.2.1.49, e.g., GH109). Degradation of cellulose and hemicellulose yields glucose and all MAGs encode glycolysis and pentose phosphate pathways (Figure 3, Supplementary Results). α-*N*-acetylgalactosaminidase releases *N*-acetylgalactosamine residues from glycoproteins that are commonly found in microbial cell walls and extracellular polysaccharides (Bodé *et al.*, 2013). *N*-acetylgalactosamine can not be directly utilized via glycolysis, however the additionally required enzymes are present (Figure 3; Supplementary Results). Under oxic conditions, organic carbon could be completely oxidized to CO_2_ via the citric acid cycle (Figure 3). Alternatively, we also identified fermentative pathways. SbA3 encodes the bifunctional aldehyde-alcohol dehydrogenase AdhE that yields ethanol (Figure 3). All MAGs encode additional aldehyde and alcohol dehydrogenases without clear substrate specificity that could also ferment acetyl-CoA to ethanol. SbA7 and SbA5 encode a L-lactate dehydrogenase (Ldh) yielding lactate from pyruvate, while six MAGs encode L-lactate dehydrogenases (LldD, GlcDEF, LutABC) that presumably perform the reverse reaction (Figure 3). Similarly, we identified pathways for acetate and / or propionate production or utilization in all MAGs (Figure 3; Supplementary Results). SbA1 and SbA3 potentially produce H_2_ via formate C-acetyltransferase PflD, which cleaves pyruvate into acetyl-CoA and formate. SbA1 encodes for the membranebound formate hydrogenlyase complex (*fdhF, hyf* operon) that produces H_2_ and might also translocate protons. SbA3 harbours an uncharacterized, cytoplasmic, monomeric FDH (*fdhA*) to transform formate to H_2_. SbA1, SbA3, SbA4, and SbA6 also encode membrane-bound, periplasmic FDH (*fdo* operon) that transfers electrons into the membrane quinol pool, as a nonfermentative alternative of formate oxidation (Figure 3, Supplementary Table S2j).

**Figure 4.**
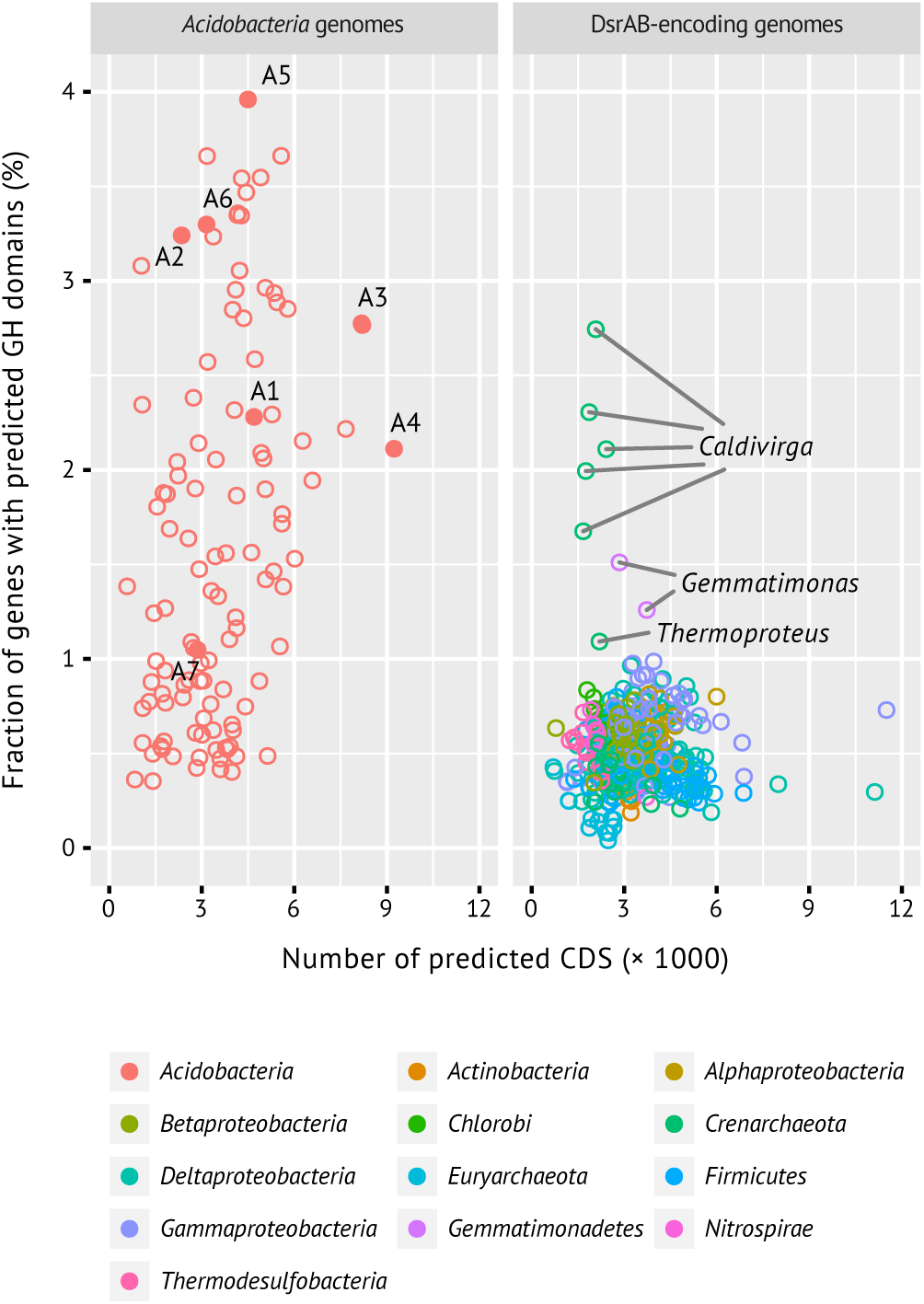
Glycoside hydrolase genes are enriched in acidobacterial genomes / MAGs compared to genomes from other taxa that encode DsrA / DsrB. DsrAB-containing MAGs SbA1 – 7 are shown as solid symbols and numbered accordingly. X-axis shows the total number of predicted CDS per genome / MAG.

### DsrAB-encoding *Acidobacteria* are metabolically active under anoxic conditions

We calculated the index of replication (iRep) with the native soil metagenomes to assess if the DsrAB-encoding *Acidobacteria* were active *in situ* (Brown *et al.*, 2016). SbA1 and SbA5, which were sufficiently complete (≥75%) for accurate measurements, had iRep values of 1.21 and 1.19, respectively. This shows that a fraction of each population was metabolically active, i.e., on average 21% of SbA1 and 19% of SbA5 cells were actively replicating at the time of sampling. Concordantly, SbA1 – 7 were also transcriptionally active in the same native soil samples. 35 – 46% of the SbA1 – 7 genes were expressed in at least one replicate. SbA1 and SbA5 contributed a considerable fraction (0.4% and 1.8%, respectively, Supplementary Table S1) of the total mRNA reads in the native soil metatranscriptome. These data likely underestimate the metabolic activity of SbA1 – 7 *in situ* because freshly sampled soil was stored at 4 °C for one week prior to nucleic acids extraction.

We further analyzed metatranscriptome data from a series of anoxic incubations of the peat soil with or without individual substrates (formate, acetate, propionate, lactate, or butyrate) and with or without supplemental sulfate (Hausmann *et al.*, 2016). While the incubations were not designed to specifically test for the MAG-inferred metabolic properties, they still allowed us to evaluate transcriptional response of the DsrAB-encoding *Acidobacteria* under various anoxic conditions (Supplementary Methods and Results). All treatments triggered shifts in genomewide gene expression; more genes were significantly (*p*<0.05) upregulated (73 – 933) than downregulated (14 – 81) as compared to the native soil. Upregulated genes included sulfur metabolism, high-affinity terminal oxidases, group 1 and 3 hydrogenases, aldehyde-alcohol dehydrogenase AdhE, glycoside hydrolases, and other carbon metabolism genes (Supplementary Table S3). Significantly upregulated glycoside hydrolase genes were GH 2, 3, 5, 9, 10, 18, 20, 23, 26, 28, 29, 30, 33, 35, 36, 38, 43, 44, 50, 51, 55, 74, 76, 78, 79, 88, 95, 97, 105, 106, 109, and 129 genes in MAGs SbA1 – 6. None of the GH genes were significantly downregulated in the incubations. Noteworthy genes that were significantly downregulated were superoxide dismutases (*sodA*) in SbA2 and SbA4 (Supplementary Table S3a).

## Discussion

Diverse members of the phylum *Acidobacteria* are abundant in various ecosystems, particularly in soils and sediments with relative abundances typically ranging from 20 – 40% (Janssen, 2006). *Acidobacteria* are currently classified in 26 subdivisions based on 16S rRNA phylogeny (Barns *et al.*, 2007). Given their phylogenetic breadth, comparably few isolates and genomes are available to explore their metabolic capabilities. Yet strains in subdivisions 1, 3, 4, and 6 are aerobic chemoorganotrophs that grow optimally at neutral or low pH (Dedysh, 2011; Eichorst *et al.*, 2011; Huber *et al.*, 2014, 2016). Furthermore, subdivision 4 contains an anoxygenic phototroph (Garcia Costas *et al.*, 2012; Tank and Bryant, 2015), subdivisions 8 and 23 contain anaerobes (Liesack *et al.*, 1994; Coates *et al.*, 1999; Losey *et al.*, 2013), subdivisions 1, 3, and 23 fermentors (Pankratov *et al.*, 2012; Kulichevskaya *et al.*, 2014; Losey *et al.*, 2013; Myers and King, 2016) and subdivision 4, 8, 10 and 23 thermophiles (Izumi *et al.*, 2012; Losey *et al.*, 2013; Crowe *et al.*, 2014; Tank and Bryant, 2015).

*Acidobacteria* have previously been described as dominant inhabitants of wetlands worldwide, namely members of subdivision 1, 3, 4, and 8 (Dedysh, 2011). Strains in the genera *Granulicella* (Pankratov and Dedysh, 2010), Telmatobacter (Pankratov *et al.*, 2012), Bryocella (Dedysh *et al.*, 2012) and *Bryobacter* (Kulichevskaya *et al.*, 2010) have been isolated from acidic wetlands and are presumably active in plant-derived polymer degradation (such as cellulose) (Dedysh, 2011; Pankratov *et al.*, 2011; Schmidt *et al.*, 2015; Juottonen *et al.*, 2017), and in nitrogen and iron cycling (Küsel *et al.*, 2008; Kulichevskaya *et al.*, 2014).

Here, we provide metagenomic and metatranscriptomic evidence that select *Acidobacteria* have a dissimilatory sulfur metabolism. The seven acidobacterial MAGs from the Schlöppnerbrunnen II peatland encode a complete dissimilatory sulfite or sulfate reduction pathway and represent novel species of at least three novel genera in subdivision 1 or 3 (Supplementary Figure S3). The phylogenetic separation into the two subdivisions in the concatenated marker gene tree is also apparent in the DsrAB tree (Supplementary Figure S1). The acidobacterial DsrAB sequences are distributed on two monophyletic clades within the uncultured family-level lineage 8 of the reductive, bacterial-type DsrAB enzyme family branch (Müller *et al.*, 2015). Furthermore, the phylogenetic breadth of the acidobacterial DsrAB sequences is representative of the intralineage sequence divergence, which suggests that the entire DsrAB lineage 8 represents yet uncultivated bacteria of the phylum *Acidobacteria*. Members of this uncultured DsrAB lineage are widespread in freshwater wetlands (Supplementary Figure S1) (Pester *et al.*, 2012) and an abundant fraction of the *dsrAB* diversity and permanent autochthonous inhabitants of oxic and anoxic soil layers in the Schlöppnerbrunnen II peatland (Steger *et al.*, 2011; Pester *et al.*, 2010).

Presence of a complete gene set for canonical dissimilatory sulfate reduction suggests that the pathway is functional, as the genetic capability for sulfate reduction can be rapidly lost by adaptive evolution if unused (Hillesland *et al.*, 2014). Except for a truncated *aprB* on SbA6, we found no indications of pseudogenes, i.e., unexpected internal stop codons or reading frame shifts, for any of the sulfate / sulfite reduction genes on the acidobacterial MAGs (Müller *et al.*, 2015). In addition, sulfur genes of each MAG were expressed in the native soil and the anoxic microcosms (Supplementary Table S3a). Many sulfur metabolism genes were also significantly upregulated in the microcosms, with *dsrC* and *aprBA* among the top 10 most expressed genes in SbA7 (Supplementary Table S3a). These findings further support full functionality of the acidobacterial dissimilatory sulfur pathways under anoxic condition.

Known SRM typically couple sulfate respiration to oxidation of fermentation products such as volatile fatty acids, alcohols, or hydrogen (Rabus *et al.*, 2013). While other microorganisms in the Schlöppnerbrunnen II soil, such as *Desulfosporosinus*, showed sulfate-and substrate-specific responses in our microcosms, hundreds of acidobacterial 16S rRNA phylotypes did not (with the exception of two) (Hausmann *et al.*, 2016). Gene expression patterns of DsrAB-encoding *Acidobacteria* in the individual anoxic microcosms as analyzed in the present study were ambiguous. Genes for putative oxidation of the supplemented substrates (formate, acetate, propionate, lactate, butyrate) were not specifically upregulated, neither with nor without supplemental sulfate. However, sulfur metabolism genes were upregulated in some incubations as compared to no-substrate-controls, suggesting indirect stimulation of sulfur dissimilation (Supplementary Results, Supplementary Table S3a). Indirect changes in microbial activity after the addition of fresh organic matter is often observed in soils (priming effects, Blagodatskaya and Kuzyakov, 2008). One priming effect explanation is the co-metabolism theory stating that easily available substrates provide the energy for microorganisms to produce extracellular enzymes to make immobile carbon accessible, which is then also available to other microorganisms. The DsrAB-encoding *Acidobacteria* have vast genetic capabilities for utilization of carbohydrates and complete sugar degradation pathways (Figure 3), in accordance with carbohydrate utilization potential previously described for subdivision 1 and 3 *Acidobacteria* (Ward *et al.*, 2009; Rawat *et al.*, 2012). Yet carbohydrate utilization coupled to sulfate reduction is a rare feature of known SRM [Cord-Ruwisch *et al.* (1986); Stetter (1988);]. While expression of many of their glycoside hydrolase genes was upregulated in our anoxic peat soil microcosms, further experiments are required to confirm that the DsrAB-encoding *Acidobacteria* couple degradation of carbohydrate polymers or monomers to sulfate reduction.

It is intriguing to propose that MAGs SbA2, SbA3, and SbA7 derive from acidobacterial SRM as they lack known sulfur oxidation genes, except *dsrL*, and express fully functional dissimilatory sulfate reduction pathways (Supplementary Table S2a), including reductive, bacterial-type *dsrAB*, and *dsrD* that may be exclusive to SRM (Hittel and Voordouw, 2000; Dahl and Friedrich, 2008; Ghosh and Dam, 2009; Rabus *et al.*, 2015). An alternative hypothesis, however, is that these novel *Acidobacteria* reverse the sulfate reduction pathway for dissimilatory sulfur oxidation or sulfur disproportionation, as was recently shown for the deltaproteobacterium *Desulfurivibrio alkaliphilus* (Thorup *et al.*, 2017). *D. alkaliphilus* also lacks known sulfur oxidation genes, except for *sqr*, and grows by coupling sulfide oxidation via an SRM-like pathway (with a reductive-type DsrAB) to the dissimilatory reduction of nitrate / nitrite to ammonium. Sulfide oxidation in acidobacterial MAGs SbA2, SbA3, and SbA7 could proceed analogous to the pathway models by Thorup *et al.* (2017) and Christiane Dahl (Dahl, 2017). Briefly, hydrogen sulfide might react with DsrC either spontaneously (Ijssennagger *et al.*, 2015) or via an unknown sulfur transfer mechanism to form persulfated DsrC. Persulfated DsrC is then oxidized by DsrMKJOP, thereby transferring electrons into the membrane quinone pool, and releasing a DsrC-trisulfide, which is the substrate for DsrAB (Santos *et al.*, 2015; Dahl, 2017). It was hypothesized that electrons released during DsrC-trisulfide oxidation to sulfite and DsrC are transferred to DsrL (Dahl, 2017). Sulfite oxidation to sulfate would be catalyzed by AprBAQmoAB and Sat.

The acidobacterial MAGs have the genomic potential to use oxygen as terminal electron acceptor and might thus couple sulfide oxidation to aerobic respiration. Alternative electron acceptors for biological sulfur oxidation in wetlands could include nitrate / nitrite and metals such as Fe(III) (Küsel *et al.*, 2008). However, known genes for dissimilatory nitrate reduction and metal reduction (Weber *et al.*, 2006) were absent from these acidobacterial MAGs. Only SbA2 encodes a putative metal reduction complex that was recently characterized in *Desulfotomaculum reducens* (Otwell *et al.*, 2015). At this time, it is unclear whether DsrAB-encoding *Acidobacteria* are capable of Fe(III) respiration, as seen in *Geothrix fermentans* (Coates *et al.*, 1999) and certain isolates in subdivision 1 (Blöthe *et al.*, 2008; Kulichevskaya *et al.*, 2014).

## Proposal of uncultivated acidobacterial genera ^*U*^*Sulfotelmatobacter*, ^*U*^*Sulfotelmatomonas*, and ^*U*^*sulfopaludibacter*

Based on combined interpretation of phylogeny (concatenated phylogenetic marker genes, DsrAB), genomic (ANI, AAI) and genetic (DsrAB) distances, and characteristic genomic features of dissimilatory sulfur metabolism (Figure 3), in accordance with Konstantinidis *et al.* (2017), we classify MAGs SbA1, SbA7, SbA5, SbA3, SbA4, and SbA6 into three new acidobacterial uncultivated genera, including uncultivated species names for the >95% complete MAGs SbA1 and SbA5. In-depth phylogenomic analysis of SbA2 was not possible and therefore it is tentatively assigned to *Acidobacteria* subdivision 3.

### *Acidobacteria* subdivision 1

- Uncultivated genus *USulfotelmatobacter* (Sul.fo.tel.ma.to.bac’ter. L. n. *sulfur*, sulfur; Gr. n. *telma, - tos*, swamp, wetland; N.L. masc. n. *bacter*, bacterium; N.L. masc. n. *Sulfotelmatobacter*, a bacterium with a dissimilatory sulfur metabolism from a swamp) with *USulfotelmatobacter kueseliae* MAG SbA1 (kue.se’li.ae. N.L. gen. n. *kueseliae*, of Kuesel, honouring Kirsten Küsel, for her work on the geomicrobiology of wetlands) and *USulfotelmatobacter* sp. MAG SbA7.
- *USulfotelmatomonas gaucii* MAG SbA5 (Sul.fo.tel.ma.to.mo.nas. L. n. *sulfur*, sulfur; Gr. n. *telma, - tos*, swamp, wetland; N.L. masc. n. *monas*, unicellular organism; N.L. masc. n. *Sulfotelmatomonas*, a bacterium with a dissimilatory sulfur metabolism from a swamp; gau’.ci.i. N.L. gen. n. *gaucii*, of Gauci, in honour of Vincent Gauci, for his pioneering work on the interplay of wetland sulfate reduction and global methane emission).

### *Acidobacteria* subdivision 3

- Uncultivated genus *USulfopaludibacter* (Sul.fo.pa.lu.di.bac’ter. L. n. *sulfur*, sulfur; L. n. *palus, - udis*, L. swamp; N.L. masc. n. *bacter*, bacterium; N.L. masc. n. *Sulfopaludibacter*, a bacterium with a dissimilatory sulfur metabolism from a swamp) with *Usulfopaludibacter* sp. MAG SbA3, *USulfopaludibacter* sp. MAG SbA4, and *USulfopaludibacter* sp. MAG S A6.
- *Acidobacteria* bacterium sp. MAG SbA2.

## Conclusion

Sulfur cycling exerts important control on organic carbon degradation and greenhouse gas production in wetlands, but knowledge about sulfur microorganisms in these globally important ecosystems is scarce (Pester *et al.*, 2012). Here, we show for the first time, using genomecentric metagenomics and metatranscriptomics, that members of the phylum *Acidobacteria* have a putative role in peatland sulfur cycling. The genomic repertoire of the novel *Acidobacteria* species encompassed recognized acidobacterial physiologies, such as facultative anaerobic metabolism, oxygen respiration, fermentation, carbohydrate degradation, and hydrogen metabolism, but was additionally augmented with a DsrAB-based dissimilatory sulfur metabolism (Figure 5). Integrating findings of sulfur oxidation in SRM and on reversibility of the dissimilatory sulfate reduction pathway (Dannenberg *et al.*, 1992; Fuseler and Cypionka, 1995; Fuseler *et al.*, 1996; Thorup *et al.*, 2017) and co-occurrence of *dsrD* and *dsrL*, genes that are considered characteristic for either sulfate reduction or sulfur oxidation (Dahl and Friedrich, 2008; Rabus *et al.*, 2015), it is conceivable that the peatland *Acidobacteria* use the same pathway for both sulfate reduction and sulfide oxidation. Some members that only encoded the pathway for dissimilatory sulfite reduction had additional genes for sulfite-producing enzymes, suggesting that organosulfonates may be a primary substrate for sulfur respiration. Our results extend our understanding of the genetic versatility and distribution of dissimilatory sulfur metabolism among recognized microbial phyla, but also underpin the challenge to unambiguously differentiate between reductive or oxidative sulfur metabolism solely based on (meta-)genome / transcriptome data (Thorup *et al.*, 2017).

**Figure 5.**
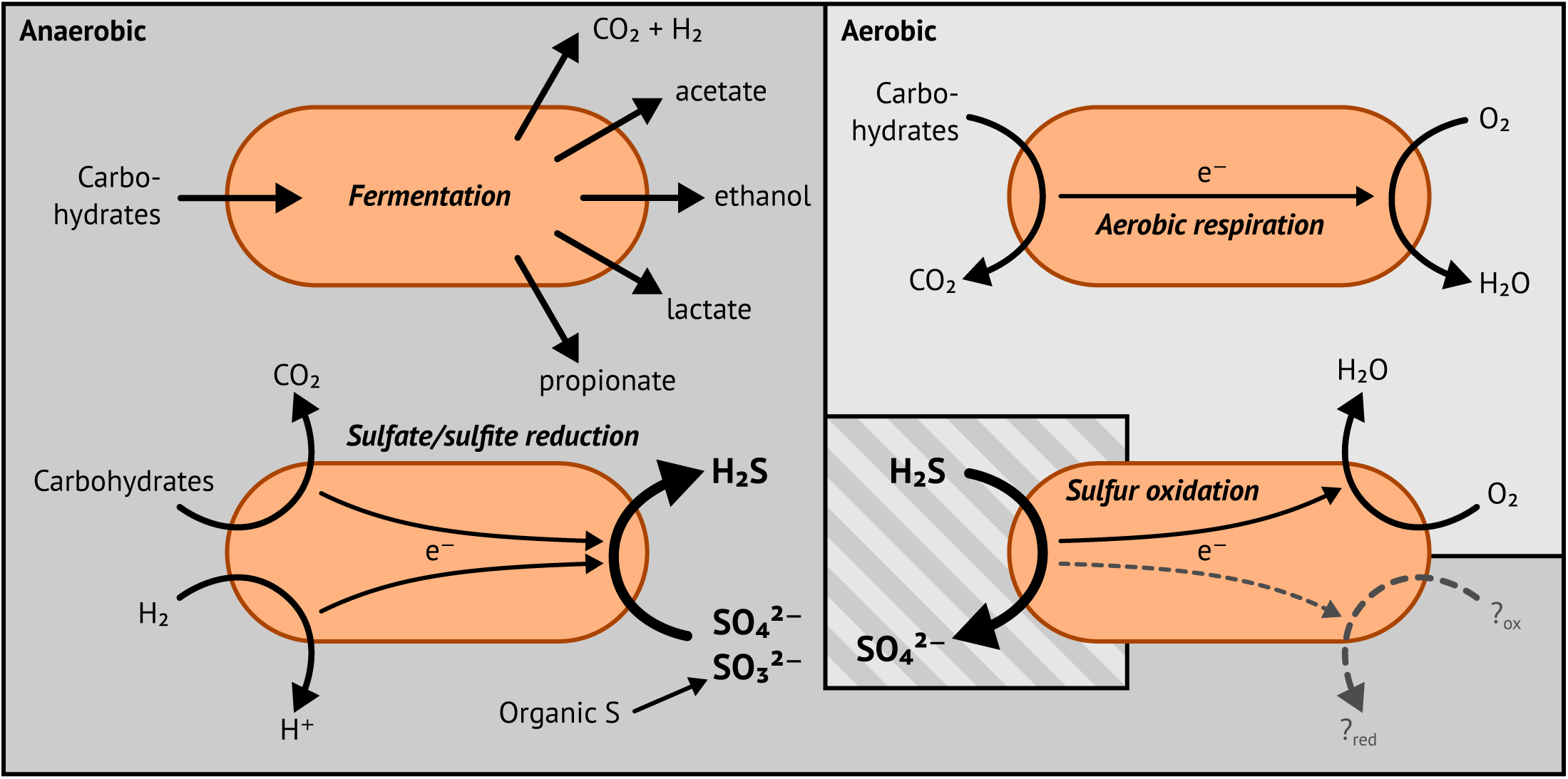
Putative lifestyles of DsrAB-encoding *Acidobacteria*.

## Acknowledgements

We are grateful to Norbert Bittner for support during field sampling, Doris Steger and Pinsurang Deevong for their contributions to qPCR analysis, and Florian Goldenberg for maintaining the Life Science Computer Cluster at the Division of Computational Systems Biology (University of Vienna). We thank the staff of the Joint Genome Institute (JGI) for metagenome and metatranscriptome library preparation, sequencing, and standard bioinformatics support, Bernhard Schink for help in naming of bacterial taxa, and Christiane Dahl, Petra Pjevac, Marc Mußmann, and Kenneth Wasmund for valuable discussions and feedback. We acknowledge LABGeM and the National Infrastructure France Genomique for providing the MicroScope MaGe platform. The work conducted by the JGI was supported by the Office of Science of the U.S. Department of Energy under Contract No. DE-AC02-05CH11231. This research was supported by the Austrian Science Fund (FWF, P23117-B17, P25111-B22, P26392-B20, and I1628-B22), the JGI (CSP 605), the German Research Foundation (DFG, PE 2147 / 1-1), and the European Union (FP7-People-2013-CIG, Grant No PCIG14-GA-2013-630188).

## Peatland *Acidobacteria* with a dissimilatory sulfur metabolism: Supplementary Information

### Supplementary Methods

#### Quantitative PCR analysis of acidobacterial subdivisions

*Acidobacteria* subdivisions 1, 2, and 3 were separately quantified using 16S rRNA gene-targeted real-time quantitative PCR (qPCR) assays. Coverage of the subdivisions was estimated using the RDP ProbeMatch online tool with RDP Release 11, Update 5, good quality filter applied, requiring a full match to the probe sequence (Cole *et al.*, 2014). The following parameters ensure optimized efficiency and sensitivity of each qPCR assay. Subdivision 1: Acid303Fa / Acid303Fb (5′-GCG CAC GGM CAC ACT GGA-3′ / 5′-GCG CGC GGC CAC ACT GGA-3′) and Acid657R (5′-ATT CCA CKC ACC TCT CCC AY-3′), 76% / 0.1% coverage by primers pairs, primer concentration: 1000 nM, annealing temperature: 68.5 °C; Subdivision 2: Acid702Fa / Acid702Fb (5′-AGA TAT CTG CAG GAA CAY CC-3′ / 5′-AGA TAT CCG CAG GAA CAT CC-3′) and Acid805R (5′-CTG ATS GTT TAG GGC TAG-3′), 64% / 7% coverage, primer concentration: 1000 nM, annealing temperature: 62.5 °C; Subdivision 3: Acid306F (5′-CAC GGC CAC ACT GGC AC-3′) and Acid493R (5′-AGT TAG CCG CAG CTK CTT CT-3′), 77% coverage, primer concentration: 500 nM, annealing temperature: 69 °C. Thermal cycling was carried out with an initial denaturation (94 °C) followed by 40 – 45 cycles of denaturation (94 °C, 40 s), annealing (68.5 °C, 62.5 °C, or 69 °C, 40 s), and elongation (72 °C, 40 – 45 s). PCR efficiency with perfectly-matched reference targets was between 82 – 86% with an R^2^ of 0.99 and a limit of detection at 100 target genes per reaction. For calculation of relative abundances, total bacterial and archeal 16S rRNA genes were quantified using a previously published qPCR assay (Pester *et al.*, 2010; Hausmann *et al.*, 2016).

#### Metagenomic and metatranscriptomic sequencing

DNA was sent to the JGI, where it was fragmented to a target length of 270 nt. Libraries were generated with the KAPA-Illumina library creation kit (KAPA biosystems) and sequenced on an Illumina HiSeq2000 sequencer. The native soil yielded 232 million 150 nt paired-end reads. Three sequencing libraries of the pooled DNA-SIP sample yielded 273, 52, and 350 million 150 nt paired-end reads each. DNA from the native soil was also sent to the King Abdullah University of Science and Technology (Thuwal, Saudi Arabia), libraries prepared with the Nextera DNA Library Prep kit (Illumina), and sequenced on an Illumina HiSeq2000 sequencer (179 million 101 nt paired-end reads).

Triplicate RNA samples from the native soil and from each incubation treatment and time point were sent to the JGI. When possible, rRNA was depleted using Ribo-Zero rRNA removal kit (Epicentre). cDNA libraries were generated with the Truseq Stranded RNA LT kit (Illumina) and sequencing was performed on an Illumina HiSeq2000 sequencer. One propionate-and sulfatestimulated replicate microcosm was excluded because of inconsistent response in sulfate turnover as compared to the other two replicates (Hausmann *et al.*, 2016), resulting in a total of 73 samples with 27 – 188 million 150 nt paired-end reads.

#### Gene-and genome-based taxonomic classification and phylogeny

Phylogenetic reconstruction of DsrAB sequences was performed based on an established DsrAB alignment (Müller *et al.*, 2015). DsrAB amino acid sequences from the MAGs and unbinned scaffolds were aligned to the filtered seed alignment using MAFFT 7.271 (Katoh and Standley, 2013). *De novo* maximum likelihood trees were calculated with FastTree 2.1.9 using the LG model and 1000 resamplings (Price *et al.*, 2010). Phylogenetic distances of the DsrAB sequences of the MAGs and scaffolds to other DsrAB harbouring organisms were calculated with T-Coffee 11 (Notredame *et al.*, 2000) using the unfiltered reference alignment without the intergenic region (Müller *et al.*, 2015). Trees were visualized with iTOL (Letunic and Bork, 2016) and Inkscape (inkscape.org).

Representative genome assemblies from the phylum *Acidobacteria* and outgroups from the *Proteobacteria, Firmicutes*, and *Verrucomicrobia* were obtained from NCBI for phylogenomic analysis. A filtered and concatenated amino acid alignment of 34 phylogenetically informative marker genes was created using CheckM (Parks *et al.*, 2015). Phylobayes was used to calculate the tree with a CAT-GTR model (Lartillot *et al.*, 2009). Phylobayes was run in five independent chains for 11000 cycles each (corresponding to approx. 6.8 × 10^6^ tree generations per chain). The first 6000 cycles in each chain were discarded as burn in (corresponding to approx. 3.7×10^6^ tree generations per chain).

Pairwise average nucleic and amino acid identities (ANI, AAI) between all protein-coding genes of each MAG and published reference genomes were calculated to estimate novelty (adapted from Varghese *et al.*, 2015). Two-way ANI and AAI were calculated based on reciprocal best blast hits filtered for sequence identity (≥70% and ≥30% for ANI and AAI, respectively) and alignment length (≥70% of the shorter sequence). Average identities and alignment fractions (AF) for each comparisons were calculated as outlined previously (Varghese *et al.*, 2015). None of the comparisons reached an ANI above the intra-species threshold of 96.5% (Varghese *et al.*, 2015). Due to lack of a generic intra-genus AAI threshold, we used the existing acidobacterial taxonomy as a reference. Intra-genus AAI variability of published acidobacterial genera with more than one species (*Acidobacterium, Granulicella, Terriglobus*) ranged from 60 – 71% (alignment fraction 52 – 66%).

#### Manually curated annotation of of DsrAB-encoding genomes

All genes of interest were manually curated using MaGe (Vallenet *et al.*, 2017). This included assessment of best-BLAST-hits to reference genomes and UniProt entries (The UniProt Consortium, 2015), presence of the required functional domains (InterPro / InterProScan; Mitchell *et al.*, 2015; Jones *et al.*, 2014) and, if appropriate, transmembrane helices (TMhmm; Krogh *et al.*, 2001), membership in the correct COGs, and membership of syntenic regions (operons). COGs in MaGe are assigned using COGnitor (www.ncbi.nlm.nih.gov/COG/) which are often too broad to be of use, therefore we additionally classified all coding DNA sequences (CDS) using the bactNOG database (eggNOG, Huerta-Cepas *et al.*, 2016). All possible HMM profiles were matched against every gene (E-value threshold 1) and only the best hit was extracted. This nonstringent E-value threshold allowed very small genes and fragments to be classified as well.

#### Carbohydrate-active enzymes

Carbohydrate-active enzymes (Lombard *et al.*, 2014, www.cazy.org) in the MAGs were identified and classified with dbCAN 4.0 (Yin *et al.*, 2012). For comparison, genomes belonging to the *Acidobacteria* and to genera with DsrAB-encoding members were downloaded from NCBI (June 2017) and analyzed with dbCAN as well. Genera with DsrAB-encoding members were identified based on literature research and UniProt InterPro / TIGRFAM searches for DsrA (IPR011806 / TIGR02064) and DsrB (IPR011808 / TIGR02066). For comparability and consistency, *de novo* ORF predictions were performed with prodigal (Hyatt *et al.*, 2010) for all genomes and MAGs. Presence of DsrA and / or DsrB was again verified with the TIGRFAM models and HMMER. dbCAN’s HMM profiles were identified with HMMER and parsed with the provided dbCAN script and R (R Core Team, 2017). Analysed acidobacterial genomes belonged to the genera *Acidobacterium, Bryobacter, Chloracidobacterium, Edaphobacter, Geothrix, Granulicella, Holophaga, Koribacter, Luteitalea, Pyrinomonas, Silvibacterium, Solibacter, Terracidiphilus, Terriglobus*, and *Thermoanaerobaculum*. 82 additional acidobacterial genomes without genus classification were also analysed. DsrA / DsrB-encoding genomes derived from the genera *Acetonema, Achromatium, Acidiferrobacter, Alkalilimnicola, Allochromatium, Ammonifex, Anaeromyxobacter, Archaeoglobus, Azospirillum, Bilophila, Caldimicrobium, Caldivirga, Carboxydothermus, Chlorobaculum, Chlorobi*um, *Curvibacter, Desulfacinum, Desulfamplus, Desulfarculus, Desulfatibacillum, Desulfatiglans, Desulfatirhabdium, Desulfatitalea, Desulfitibacter, Desulfitobacterium, Desulfobacca, Desulfobacter, Desulfobacterium, Desulfobacula, Desulfobulbus, Desulfocapsa, Desulfocarbo, Desulfococcus, Desulfocurvus, Desulfofervidus, Desulfofustis, Desulfohalobium, Desulfoluna, Desulfomicrobium, Desulfomonile, Desulfonatronospira, Desulfonatronovibrio, Desulfonatronum, Desulfonauticus, Desulfonispora, Desulfopertinax, Desulfopila, Desulfoplanes, Desulforegula, Desulforhopalus, Desulforudis, Desulfosarcina, Desulfospira, Desulfosporosinus, Desulfotalea, Desulfothermus, Desulfotignum, Desulfotomaculum, Desulfovermiculus, Desulfovibrio, Desulfovirgula, Desulfurella, Desulfurispora, Desulfurivibrio, Dethiosulfatarculus, Dissulfuribacter, Ferriphaselus, Gallionella, Gemmatimonas, Gordonibacter, Gracilibacter, Halodesulfovibrio, Halorhodospira, Lamprocystis, Lautropia, Magnetococcus, Magnetomorum, Magnetoovum, Magnetospira, Magnetospirillum, Magnetovibrio, Marichromatium, Moorella, Pelodictyon, Phaeospirillum, Prosthecochloris, Pyrobaculum, Rhodomicrobium, Rubrivivax, Ruegeria, Ruthia, Sedimenticola, Sideroxydans, Sulfuricella, Sulfuritalea, Syntrophobacter, Syntrophomonas, Syntrophus, Thermanaeromonas, Thermocladium, Thermodesulfatator, Thermodesulfobacterium, Thermodesulfobium, Thermodesulforhabdus, Thermodesulfovibrio, Thermoproteus, Thermosinus, Thermosulfurimonas, Thioalkalivibrio, Thiobacillus, Thiocapsa, Thiocystis, Thiodiazotropha, Thioflavicoccus, Thioflexothrix, Thioglobus, Thiohalocapsa, Thiohalomonas, Thiolapillus, Thiomargarita, Thioploca, Thiorhodococcus, Thiorhodovibrio, Thiosymbion, Thiothrix*, and *Vulcanisaeta*.

#### Expression analysis

Metatranscriptomic reads were quality filtered at the JGI using their analysis pipeline. In short, the raw reads were quality-trimmed to Q10, adapter-trimmed using bbduk (minimal allowed length 50 nt), followed by removal of PhiX control sequences, artefacts, human sequences, and reads containing N bases with bbduk / bbmap (BBTools, http://jgi.doe.gov/data-andtools/bbtools/). rRNA reads were removed using the SILVA database (Quast *et al.*, 2013) and bbmap. This resulted in 73 samples with 22 – 161 million high quality non-rRNA reads with a median length of 150 nt. Those were then mapped to the combined metagenomic assembly using bowtie2 with the default scoring function (Langmead and Salzberg, 2012). Fragments per CDS were then independently counted using featureCounts 1.5.0 (Liao *et al.*, 2014). Differential expression analysis of SbA1 – 7 was performed using R (R Core Team, 2017) and the DESeq2 package (Love *et al.*, 2014).

### Supplementary Results and Discussion

#### Sulfite dehydrogenase homologs

We identified several genes encoding putative sulfite dehydrogenases of the COG2041 family: (A) SbA3 and SbA4 encode orthologs to *Cupriavidus necator* (Ralstonia eutropha) N-1 *soxC* (CNE_1c35220), Sbm_v1_d1920025 and Sbm_v1_e3730002, respectively (∽50% sequence identity). Directly downstream are genes homologous to the N-terminal region of *C. necator soxD* (CNE_1c35210), Sbm_v1_d1920026 and Sbm_v1_e3730003, respectively (∽40% sequence identity at <50% overlap). *soxABXYZ*, present in *C. necator*, are not found on any acidobacterial MAG. However, *C. necator* N-1 can not oxidize thiosulfate (Dahl and Friedrich, 2008). (B) *Ca. Solibacter usitatus* encodes a *sorAB*-like gene pair (Acid_7248 – 7249), which we also found in SbA1 (Sbm_v1_b530003 – 4), SbA3 (Sbm_v1_d50016 – 17), SbA4 (Sbm_v1_e5130014 – 13 and fragmented Sbm_v1_e7680001 – 2), and SbA6 (Sbm_v1_g110054 – 55). The *sorAB*-like genes from the MAGs are <40% identical to Starkeya novella SorAB (Snov_3268 – 3269). SorA transfers electrons from oxidizing sulfite to the membrane-bound cytochrome c SorB subunit. SorB is then oxidized by a terminal oxidase. SorAB in S. novella is potentially involved in aerobic respiration or in sulfite detoxification (Simon and Kroneck, 2013). (C) YedYZ-like proteins are present in SbA1 (Sbm_v1_b880028 – 29), SbA4 (Sbm_v1_e240001 – 2, fragmented), and SbA5 (Sbm_v1_f120015 – 16), which are related to the sulfide oxidase family and are part of COG2041. Their function is unknown (Dahl and Friedrich, 2008) and they are found in several other acidobacterial genomes and *E. coli*. (D) Additional, completely uncharacterized members of COG2041 are present in SbA3, SbA4, and SbA5.

#### Respiration and oxidative stress

Respiration with oxygen as terminal electron acceptor requires a membrane electron transport chain involving up to four complexes, i.e., the NADH dehydrogenase (NDH, respiratory complex I), the succinate dehydrogenase (SDH, respiratory complex II), the quinol — cytochrome-c reductase (cytochrome *bc*_*1*_ complex or alternative complex III, respiratory complex III), and the terminal oxidase (cytochrome-c oxidase or cytochrome *bd*-type oxidase, respiratory complex IV). Complexes I (NDH-1 only), III, and IV (except *bd*-type) translocate protons through the cell membrane building up proton motive force. The ATP synthase uses the proton motive force to generate ATP and is called respiratory complex V. SbA5 and SbA7 encode every gene for complexes I – V, while SbA1, SbA3, SbA4, and SbA6 encode only partial operons for some complexes. SbA2, the most incomplete MAG, is lacking all genes of complex II and V (Figure 3).

Oxidation of organic matter generates reducing equivalents, e.g., in glycolysis one NADH is formed per one glucose. NADH is oxidized to NAD^+^/H^+^ by the NDH, which in turn transfers electron to the membrane quinone pool. Two types of NDH are characterized in *E. coli*. NDH-1, consisting of a large complex encoded by the *nuo* operon, translocates protons, while NDH-2, encoded by a single gene (*ndh*), does not. We identified both NDH-1 and NDH-2 in the MAGs, with the latter (partially) missing in SbA2, SbA4, and SbA7. All MAGs harbour one or more (partially fragmented) *nuoACDHJKLMN* operons. One operon in each MAG also includes *nuoEFG* (except in SbA3). NuoEFG forms the catalytic NADH dehydrogenase module of complex I, while NuoBCDHIN and NuoKLM form the hydrogenase and transporter modules, respectively. NuoA and NuoJ are not part of the modules and likely involved in assembly of the complex (Friedrich *et al.*, 2016). SbA6 is missing *nuoI*, likely because of MAG incompleteness (Supplementary Table S2b).

SDH is encoded by the *sdh* operon. It consists of a cytoplasmic-facing catalytic subunit (SdhAB) and a transmembrane cytochrome b_*556*_ or b_*558*_ subunit (SdhCD in *E. coli* or larger SdhC in B. subtilis), which together transfer the electrons from oxidation of succinate to fumarate into the membrane quinone pool. It is the only complex of the respiration chain not involved in proton translocation. Both complex I and II are found in many anaerobically respiring microorganisms, including SRM (e.g., Pereira *et al.*, 2011; Klenk *et al.*, 1997; Rabus *et al.*, 2004; Strittmatter *et al.*, 2009; Plugge *et al.*, 2012; Visser *et al.*, 2013; Kuever *et al.*, 2014; Mardanov *et al.*, 2016). We identified SDH in all MAGs but SbA2, arranged like the B. subtilis-type operon (*sdhCAB*) (Supplementary Table S2c). SbA1 harbours a second operon (*sdhACDB*) that could also be a fumarate reductase (*frdACDB*). Fumarate reductase performs the reverse reaction of SDH as part of the reductive citric acid cycle, but both enzymes were shown to catalyze both reaction in *E. coli* (Guest, 1981; Maklashina *et al.*, 1998). No other MAG harboured candidates for fumarate reductase.

Complex III, the quinol — cytochrome-c reductase transfers electron from the membrane quinone pool to cytochrome c. Cytochrome c is then oxidized and O2 is reduced to H_2_O by a cytochromec terminal oxidase. Alternately, a quinol terminal oxidase can directly utilize electrons from the membrane quinone pool to reduce O_2_. Two isofunctional complexes of quinol — cytochrome-c reductases are known. The two component cytochrome *bc1* complex is encoded by the *pet* operon and present in all MAGs but SbA2 (Supplementary Table S2d). Alternative complex III (ACIII) (Refojo *et al.*, 2012) with seven subunits is encoded by the *act* operon and present in all MAGs but SbA7 (Supplementary Table S2e). Some of the terminal oxidase genes are found immediately down-or upstream of quinol — cytochrome-c reductase operons (Supplementary Tables S2e – g), as is observed in other *Acidobacteria* (e.g., *Ca. K. versatilis, Ca. S. usitatus, Chloracidobacterium thermophilum*; Garcia Costas *et al.*, 2012). In between alternative complex III and the terminal oxidase genes, we always find the *sco* gene coding for a chaperone of the SCO1 / SenC family, a putative assembly factor for respiratory complexes (Buggy and Bauer, 1995). We identified both families of terminal oxidases, i.e., haem-copper oxidases (HCO) of classes A and C, which are cytochrome-c oxidases, and cytochrome *bd*-type oxidases, which are quinol oxidases. HCO class A are classified as low-affinity terminal oxidases (LATO) in contrast to HCO class C and *bd*-type oxidases that are high-affinity terminal oxidases (HATO) (Morris and Schmidt, 2013). Three distinct operon structures for HCO family A were observed in SbA1 – 7 and also other *Acidobacteria*, e.g. *Ca. K. versatilis* and *Ca. S. usitatus*: (1) downstream of ACIII and *sco*, (2) upstream of *petBC* (not found in *Ca. K. versatilis*), both named *ctaCDEF*, and (3) without quinol — cytochrome-c reductase genes found up-or downstream but with a subunit III consisting of two genes (*coxOP*) (Supplementary Table S2f). The subunit II of all three types have the CuA copper center motifs (IPR001505), which is found in cytochrome-c oxidases but not in quinol oxidases (Pereira *et al.*, 2001). The essential subunits are I and II (Pereira *et al.*, 2001) and are present in all MAGs but SbA4. High-affinity terminal oxidases of HCO class C or *bd*-type oxidases are present in SbA1, SbA5, SbA3, and SbA6 (Supplementary Table S2g). Both types are encoded by two subunits on the MAGs. Secondary genes, as found in other organisms, e.g., *ccoQP* (Bühler *et al.*, 2010) or *cydS* / *cydX* (Cook and Poole, 2016), are missing.

An ATP synthase of the F_o_F_1_-ATPase type is present in all MAGs but SbA2 (Supplementary Table S2h). Its genes are consistently split into two operons, *atpZIBE* and *atpF*′FHAGDC. The former is missing in SbA6, while the later is fragmented in SbA1 with *atpG* missing completely. *atpF* and *atpF*′ are paralogs of the subunit B. In cyanobacteria, a homodimer of subunit B is replaced by a heterodimer of subunit B and B′ (Dunn *et al.*, 2001). However, *atpF*′F is found in nonphotosynthetic *Acidobacteria*, e.g., *Ca. K. versatilis* and *Ca. S. usitatus*, and also in the photoheterotroph *Chloracidobacterium thermophilum*.

A second function is attributed to terminal oxidases in some organisms, i.e., defence against oxidative stress, especially to *bd*-type oxidases (Giuffrè *et al.*, 2014). The strictly anaerobic SRM *Desulfovibrio* vulgaris encodes two terminal oxidases, one cytochrome-c oxidase (HCO class A, DVU_1815 – 1812) and one *bd*-type oxidase (DVU_3271 – 3270). It was demonstrated that both are involved in the detoxification of oxygen (Ramel *et al.*, 2013). Terminal oxidases are also needed to remove oxygen produced by superoxide detoxification. Superoxide dismutase converts superoxide to oxygen and hydrogen peroxide. Hydrogen peroxide is then removed by catalases, peroxidases, or glutathione peroxidases (Figure 3). Manganese-dependent superoxide dismutase (*sodA*) is found in all MAGs and Cu-Zn-dependent superoxide dismutase (*sodC*) in SbA2 and SbA7. SbA1, SbA2, SbA3, and SbA5 encode for bifunctional haem-dependent catalaseperoxidases (*katG*), but none of the seven MAGs for mono-functional, haem-dependent catalases (*katE* / *katA*) or manganese-dependent catalases (*katN*). SbA2 and SbA4 encode for glutathione peroxidases (Supplementary Table S2i).

**Dissimilatory nitrogen metabolism and nitrogen fixation**

Although nitrate availability is limited in wetlands (Pester *et al.*, 2012a), we investigated the possibility for nitrate respiration in the MAGs. SRM contribute to nitrogen cycling, as some can respire nitrate / nitrite as an alternative electron acceptor to sulfate or fix atmospheric nitrogen using nitrogenase (Rabus *et al.*, 2013; Marietou, 2016). Oxidation of sulfur compounds coupled to nitrate or nitrite reduction is common among sulfide-/ thiosulfate-oxidizing microorganisms (Ghosh and Dam, 2009), but was also observed in organisms encoding reductive-type DsrAB genes. *Desulfovibrio desulfuricans, Desulfobulbus propionicus*, and *Desulfurivibrio alkaliphilus* were shown to oxidize sulfide with nitrate / nitrite as the electron acceptor (Dannenberg *et al.*, 1992; Thorup *et al.*, 2017). It is also proposed that the sulfide-oxidizing cable bacteria (*Desulfobulbaceae*) can use nitrate / nitrate as an alternative to oxygen (Marzocchi *et al.*, 2014).

Only few *Acidobacteria* were shown to perform nitrate reduction or encode the required marker genes (e.g., Ward *et al.*, 2009; Männistö *et al.*, 2012). SbA1 – 7 lack *narGHI, napAB, nrfA, nirK, nirS, norBC*, and *nosZ* (Kraft *et al.*, 2011) and thus the genomic potential for dissimilatory nitrate or nitrite reduction. Only SbA5 harbours two gene copies of the nitric oxide reductase NorZ (also known as qNOR), an enzyme that is likely not involved in denitrification but used for nitric oxide detoxification (Kraft *et al.*, 2011). The MAGs also lacked key genes of aerobic nitrogen metabolisms i.e., *amoCAB* (ammonia oxidation), *nxrAB* (nitrite oxidation), and *nifH* (nitrogen fixation) (Pester *et al.*, 2012b, 2014; Gaby and Buckley, 2014; Daims *et al.*, 2016).

#### Dissimilatory metal reduction

The genes required for dissimilatory metal reduction (*mtr / omc* operon) as described for *Shewanella* and *Geobacter* (Shi *et al.*, 2006; Weber *et al.*, 2006; Coursolle and Gralnick, 2010) are absent in all MAGs. Direct interspecies electron transfer (DIET) is an important but understudied process in wetlands (Holmes *et al.*, 2017). However, we could not identify any homologs to the essential pilin-associated c-type cytochrome OmcS (Shrestha *et al.*, 2013; Holmes *et al.*, 2017). We found an ortholog to a novel metal reduction complex in *Desulfotomaculum reducens* (Dred_1685 – 1686) (Otwell *et al.*, 2015) in SbA2 (Sbm_v1_c100009 – 10). This complex was shown to reduce Fe(III), Cr(VI), and U(VI) with NADH as the electron donor, but its physiological role is unresolved (Otwell *et al.*, 2015).

#### Import and phosphorylation of glucose

All genomes except SbA7 harbour at least one cytoplasmic glucokinase (*glk* / *glcK*) (Supplementary Table S2m). Cytoplasmic glucokinases are required to utilize glucose released by cytoplasmic polysaccharides degradation, but are not required for growth on glucose, as extracellular glucose can be imported and phosphorylated by the phosphotransferase system (PTS). The PTS is missing in all MAGs — Enzyme I and histidine protein are both not present. Only one fragment of a mannitol-specific enzyme IIBC component is found in SbA2 (Sbm_v1_c130007). Alternately, Lindner *et al.* (2011) demonstrated that inositol permeases IolT1 / IolT2 are low-affinity glucose permeases, and together with glucokinases can replace the PTS in Corynebacterium glutamicum. *iolT1* and *iolT2* match TIGRFAM’s sugar porter motif TIGR00879, which we also find in genes in all genomes but SbA2 (data not shown). These could putatively transport glucose, but also other sugars or inositol.

#### *N*-acetylgalactosamine degradation

*N*-acetylgalactosamine (GalNAc) degradation consists of five steps before entering glycolysis: (1) *N*-acetylgalactosamine kinase, (2) *N*-acetylgalactosamine-6-phosphate deacetylase, (3) galactosamine-6-phosphate deaminase, (4) tagatose-6-phosphate kinase, and (5) tagatosebisphosphate aldolase (Figure 3, Supplementary Table S2l). Neither GalNAc-specific PTS genesor *N*-acetylgalactosamine kinase (*agaK*) were identified. However, other sugar kinases of are resent, e.g., *glcK*-type glucokinases. Not all sugar kinases are specific for only one substrate (e.g., Reith *et al.*, 2011), therefore some of the those could putatively act as *N*acetylgalactosamine kinases. *N*-acetylglucosamine-6-phosphate deacetylase (*nagA*) was found in six MAGs. *E. coli* possesses two homologous *N*-acetylglucosamine-6-phosphate deacetylases (*nagA* and *agaA*, COG1820), both of whom can utilize GalNAc and *N*-acetylglucosamine (GlcNAc) (Hu *et al.*, 2013). Thereby it is likely the identified genes code for bifunctional deacetylases as well. Galactosamine-6-phosphate deaminase (AgaS), found in SbA1, SbA7, and SbA6, converts D-galactosamine 6-phosphate to D-tagatofuranose 6-phosphate. PfkB, found in SbA3, SbA4, and SbA2 is a bifunctional 6-phosphofructokinase and tagatose-6-phosphate kinase in *E. coli* (Babul, 1978). Most MAGs contain the last enzyme needed, tagatose-bisphosphate aldolase (LacD), which is a class I aldolase that produces glycerone phosphate and D-glyceraldehyde 3-phosphate. The *E. coli* class II aldolases of the same function (*kbaYZ, gatYZ*) are heteromeric. Only the noncatalytic subunit *kbaZ* is found in SbA1 (Sbm_v1_b1710004). *kbaY* was never found.

#### Lactate, propionate, and butyrate metabolism

Six of the acidobacterial MAGs harbour four different types of L-lactate dehydrogenases, while SbA6 has none (Supplementary Table S2o). NAD-dependent L-lactate dehydrogenase Ldh (SbA7 and SbA5) ferments pyruvate to L-lactate anaerobically, while the FMN-dependent L-lactate dehydrogenase LldD (SbA1, SbA7, SbA3, and SbA4), L-lactate / D-lactate / glycolate dehydrogenase GlcDEF (SbA5, SbA3, and SbA4) and, LUD-type L-lactate dehydrogenase LutABC (SbA5, SbA2, SbA3, and SbA4) utilize L-lactate as an energy and carbon source. We found putative D-lactate dehydrogenases (Dld) (SbA1, SbA5, and SbA2), which probably convert Dlactate to pyruvate. These are homologs to *Archaeoglobus* fulgidus Dld (AF_0394) and mitochondrial D-lactate dehydrogenases (<30% identity).

With the exception of SbA2, all MAGs contain key genes for propionate oxidation with complete pathways found in SbA1 and SbA5 (Supplementary Table S2o). Conversion of propionate to propionyl-CoA is performed by a CoA transferase. Propionyl-CoA:succinate CoA transferase ScpC is encoded in SbA1 and acetate CoA-transferase YdiF, a family 1 CoA transferase, which has propionyl-CoA:acetate CoA transferase activity (Rangarajan *et al.*, 2005), is encoded in SbA5 and SbA4 (Supplementary Table S2o). Other family 1 CoA transferase genes (IPR004165) are present,except in SbA2. It is unclear if these can utilize propionate as well, however it was proposed before for *Desulfotomaculum kuznetsovii* (Visser *et al.*, 2013). The main subunit gene of the propionyl-CoA carboxylase (PccB), which produces (*S*)-methylmalonyl-CoA, is present in all six MAGs. The propionyl-CoA carboxylase biotin carboxylase subunit and biotin carboxyl carrier protein are, however, missing in SbA7 and SbA4. Stereochemical inversion to (*R*)-methylmalonyl-CoA is performed by methylmalonyl-CoA epimerase (Mce), which is encoded in the same MAGs as ScpC / YdiF. Methylmalonyl-CoA mutase, catalyzing the final step in the pathway, is present in all six MAGs (Supplementary Table S2o).

Various putative beta-oxidation genes are present in all MAGs, however the substrate specificities of their encoded enzymes are unclear. For the case of butyrate oxidation, the physiological mechanisms are resolved in detail in the syntrophic organism *Syntrophomonas wolfei* (Schmidt *et al.*, 2013). No orthologs to the key enzyme butyryl-CoA dehydrogenase (Swol_1933 / Swol_2052) are present in any of the MAGs, therefore it is unlikely they can perform (syntrophic) butyrate oxidation.

#### Differential gene expression between anoxic microcosm incubations

To analyze changes in gene expression of SbA1 – 7 during the anoxic peat soil incubations, we performed pairwise comparisons between different treatments and time points: (1) at every time point and for each added substrate we compared the microcosms amended with sulfate to those without external sulfate (i.e., stimulation or downregulation caused by sulfate), (2) at every time point and separately for sulfate-stimulated incubations and no-sulfate-controls we compared microcosms amended with substrate to the no-substrate-controls (i.e., up-or downregulation caused by formate, acetate, propionate, lactate, or butyrate); and (3) for each treatment we compared the early time point (8 days) to the late time point (36 days) (Supplementary Table S3a). When compared to the gene expression changes between the native soil and the incubations, less genes were upregulated between different incubations treatments. Differentially expressed genes included dissimilatory sulfur genes, hydrogen metabolism genes, electron transfer genes, and a few genes belonging to the tricarboxylic acid cycle (Supplementary Table S3a).

Compared to the butyrate-only incubation, expression of *sat, aprBA, qmoBC, dsrAB, dsrN, dsrT*, and *srL* of SbA2 was induced upon addition of sulfate and butyrate. One subunit of (2*R*)-sulfolactate sulfo-lyase (*suyB*) was overexpressed in SbA4 in incubations with sulfate and formate. Compared to the no-substrate-controls, we observed significant overexpression of some sulfur metabolism genes from SbA2, SbA3, and SbA7 in formate-, propionate-, lactate-, and / or butyrate-amended incubations, with and / or without supplemental sulfate (Supplementary Table S3a). However, we observed no significant expression changes in the genes that are possibly involved in oxidation of the amended substrates. Moderate expression of sulfate reduction genes without addition of external sulfate is expected due to cryptic sulfur cycling under anoxic conditions (Pester *et al.*, 2012a). The peat soil microcosms without external sulfate contained low amounts of endogenous sulfate (24 ± 6 μM) that was only depleted after 11 – 25 days of incubation (Hausmann *et al.*, 2016).

Group 3 hydrogenase gene were affected by substrate amendment in incubation with and / or without external sulfate added. Group 3b hydrogenase (*hyhBCSL*) of SbA2 was stimulated by all substrates except acetate. Group 3c hydrogenase (*mvhDCA*) of SbA4 was significantly downregulated after the addition of propionate, lactate, and butyrate. Group 3d hydrogenase (*hoxEFYH*) of SbA1 was significantly overexpressed in butyrate-amended microcosms. HoxF of SbA2 was overexpressed when lactate or butyrate was added. Group 3 hydrogenases are cytoplasmic and possibly bidirectional (Greening *et al.*, 2016), leaving it unresolved if hydrogen is produced from the substrates or if hydrogen is provided from a substrate-utilizing syntrophic partner.

## Supplementary Tables

**Supplementary Table S1**

Taxonomy, genome characteristics, and abundance measures of the DsrAB-encoding MAGs. Estimation of completeness and contamination was performed with checkM. Genome abundance estimates are the fraction of metagenomic reads mapped to each MAG in relation to all quality filtered reads. Values given for native soil are averages of both native soil metagenomes. mRNA abundance estimates (only acidobacterial MAGs) are the fraction of fragments (paired-end reads) mapped to all of each MAG’s CDS in relation to all quality filtered fragments. Standard deviation of three replicates is given. Fraction of expressed CDS in native soil (%) is given only for acidobacterial MAGs.

**Supplementary Table S2**

Curated annotation tables of SbA1 – 7: (a) dissimilatory sulfur metabolism; (b – h) respiratory complexes I – V: NADH dehydrogenases (NDH, b), succinate dehydrogenase (SDH, c), quinol — cytochrome-c reductases (CIII / ACIII, d / e), low-(LATO, f) and high-affinity (HATO, g) terminal oxidases, and ATP synthases (h); (i) stress (superoxide detoxification), (j) formate dehydrogenases (FDH), hydrogenases (Hase); (k) cytoplasmic electron transport systems; (l) *N*acetylgalactosamine degradation; (m) glycolysis (and gluconeogenesis), pentose phosphate pathway, and Entner-Doudoroff pathway; (n) citric acid cycle (TCA); (o) pyruvate, acetate, propionate, and related metabolisms; (p) dissimilatory metal metabolism. Columns provide functional categories (only a, m, o), pathway step number and / or proposed direction (only m, o), product (enzyme, transporter) names with EC and TC numbers where appropriate, subunit names or descriptions where appropriate, gene names, and locus numbers per MAG. All loci are prefixed by Sbm_v1_. Products with multiple copies per MAG are separated into more than one column (only b – f, j). Fragmented genes (assembly or biological artefacts) are marked with downward arrows (↓) after their loci numbers. TM, transmembrane subunit. ^1^ or ^2^ indicates the first or last CDS on a scaffold (depending on the reading frame).

The following metabolic marker genes are absent in SbA1 – 7 MAGs: Inorganic sulfur metabolism (Wasmund *et al.*, 2017): *tsdA*, thiosulfate dehydrogenase; *otr*, octaheme tetrathionate reductase; *phsABC*, thiosulfate reductase; *psrABC*, polysulfide reductase; *sreABC*, sulfur reductase (Laska *et al.*, 2003); *asrABC*, siroheme-independent dissimilatory sulfite reductase (anaerobic sulfite reductase) (Huang and Barrett, 1991); *fsr*, coenzyme F_420_-dependent sulfite reductase (Johnson and Mukhopadhyay, 2005). ATPases: *atpDCQRBEFAG*, N-ATPase (alternative “archaeal-type” F?F1-ATPase) (Sumi *et al.*, 1997; Dibrova *et al.*, 2010). Dissimilatory nitrate reduction to ammonium (DNRA) (Kraft *et al.*, 2011): *napAB*, periplasmic nitrate reductase (NAP), catalytic subunit is NapA; *nrfA*, periplasmic cytochrome c nitrite reductase (catalytic subunit). Denitrification (Kraft *et al.*, 2011): *narGHI*, membrane bound cytoplasm-facing nitrate reductase (NAR), catalytic subunit is NarG; *nirK* or *nirS*, isofunctional but evolutionarily unrelated periplasmic nitric oxide-forming nitrite reductase (NIR); *norBC*, membrane bound periplasmfacing nitric oxide reductase (NOR), catalytic subunit is NorB; *nosZ*, periplasmic nitrous oxide reductase (NOS). Nitrogen fixation: *nifH*, nitrogenase. Nitrification: *amoCAB*, ammonia monooxygenase (AMO); *nxrAB*, nitrite oxidoreductase (NXR). Methanotrophy (Iguchi *et al.*, 2010): *pmoCAB*, particulate methane monooxygenase (pMMO); *mmoXYBZDC*, soluble methane monooxygenase (sMMO). Photosynthesis: *pscAB*-*fmoA, Chloracidobacterium thermophilum* photosystem (Garcia Costas *et al.*, 2012); *pscABCD, Chlorobium tepidum* photosystem (Eisen *et al.*, 2002); *pufABCMLH, Allochromatium vinosum* photosystem (Weissgerber *et al.*, 2011). ROS defence: *katN*, mono-functional, manganese catalase (EC 1.11.1.6) (Wu *et al.*, 2004); *katE* / *katA*, mono-functional, haem-containing catalase (EC 1.11.1.6). Hydrogenases: [FeFe] hydrogenases.

**Supplementary Table S3**

(a) Supplementary Table S2 deposited in machine-readable format including additional information. The length of CDS terminated by scaffold borders (^1^ or ^2^ in strand column) is underestimated, as the true length is not known. bactNOG and NOG IDs were assigned by bestmatch principle. Ranks are based on averaged FPKM. Missing ranks indicate that expression was never detected in any replicate. Significant differential expression is shown separated by factor. Three-letter-codes are initials of amended substrates (F, A, P, L, B) or N for no-substrate-control, sulfate stimulation (S) or control without external sulfate (C), and early (E, 8 days) and late (L, 36 days) time points, followed by the log2 fold change. Over-or underexpression is indicated by arrows of small / larger than signs. (b) Glycoside hydrolase genes identified with dbCAN.

**Supplementary Table S4**

Glycoside hydrolases genes summarized by EC numbers provided by the carbohydrate-active enzymes database.

## Supplementary Figures

**Supplementary Figure S1.**
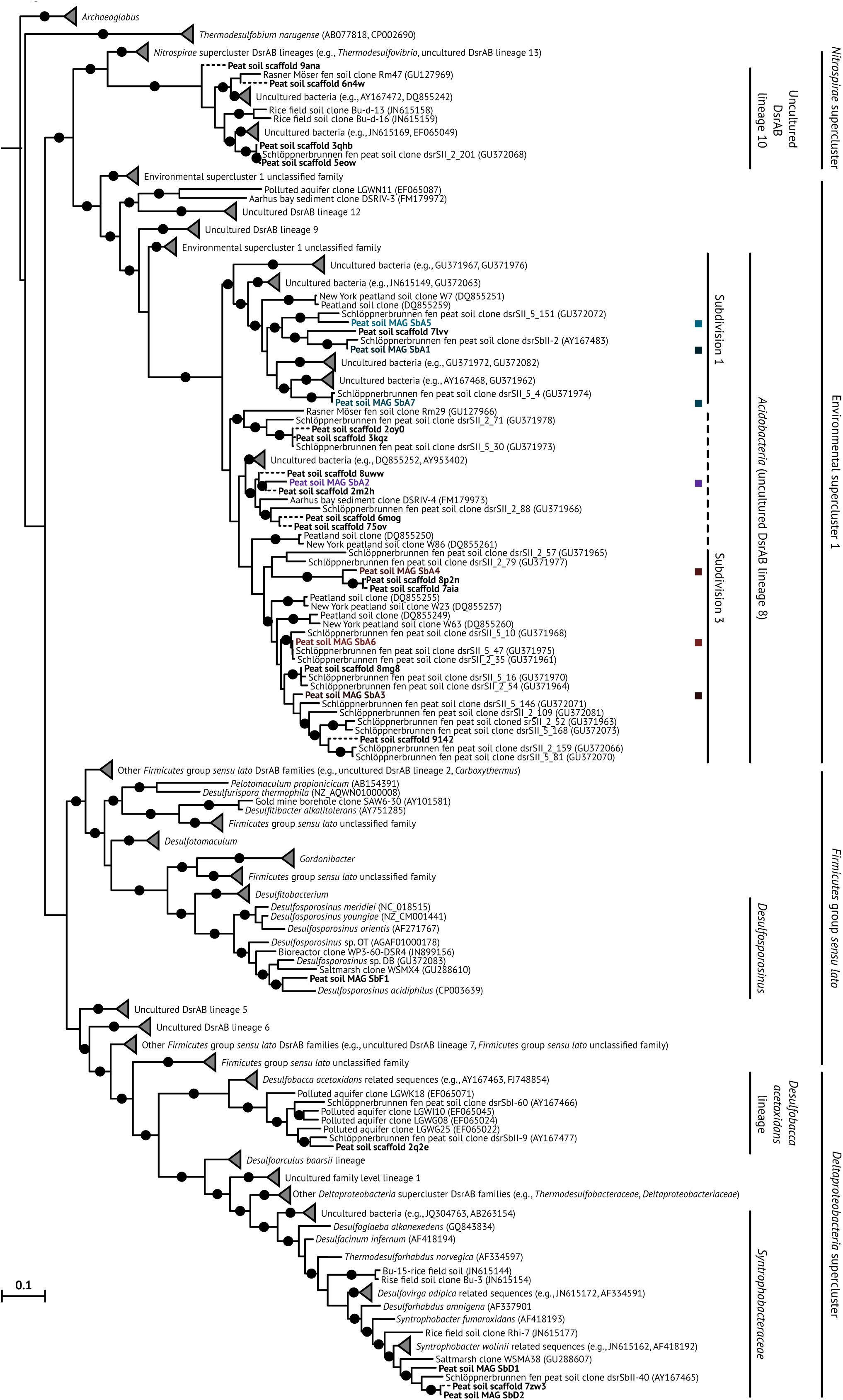
Reductive bacterial-type DsrAB. Maximum likelihood tree was calculated by FastTree (LG model, 1000 resamplings) using a reference amino acid alignment with reductive bacterial-type DsrAB indel positions removed (Müller *et al.*, 2015). Branch supports equal to or greater than 0.9 are indicated by black circles. DsrAB sequences from MAGs and scaffolds are marked in bold. Binned acidobacterial DsrAB sequences are coloured analogous to Figure 3. Dashed branches represent incomplete *dsrAB* gene sequences (e.g., caused by a contig ending) that were sufficiently long to be included in the phylogenetic analysis (only *dsrAB* genes on scaffold 43ik were too short and omitted). The extent of subdivision 3 group is unclear and indicated by a dashed line. Outgroup sequences are shown in Supplementary Figure S2.

**Supplementary Figure S2.**
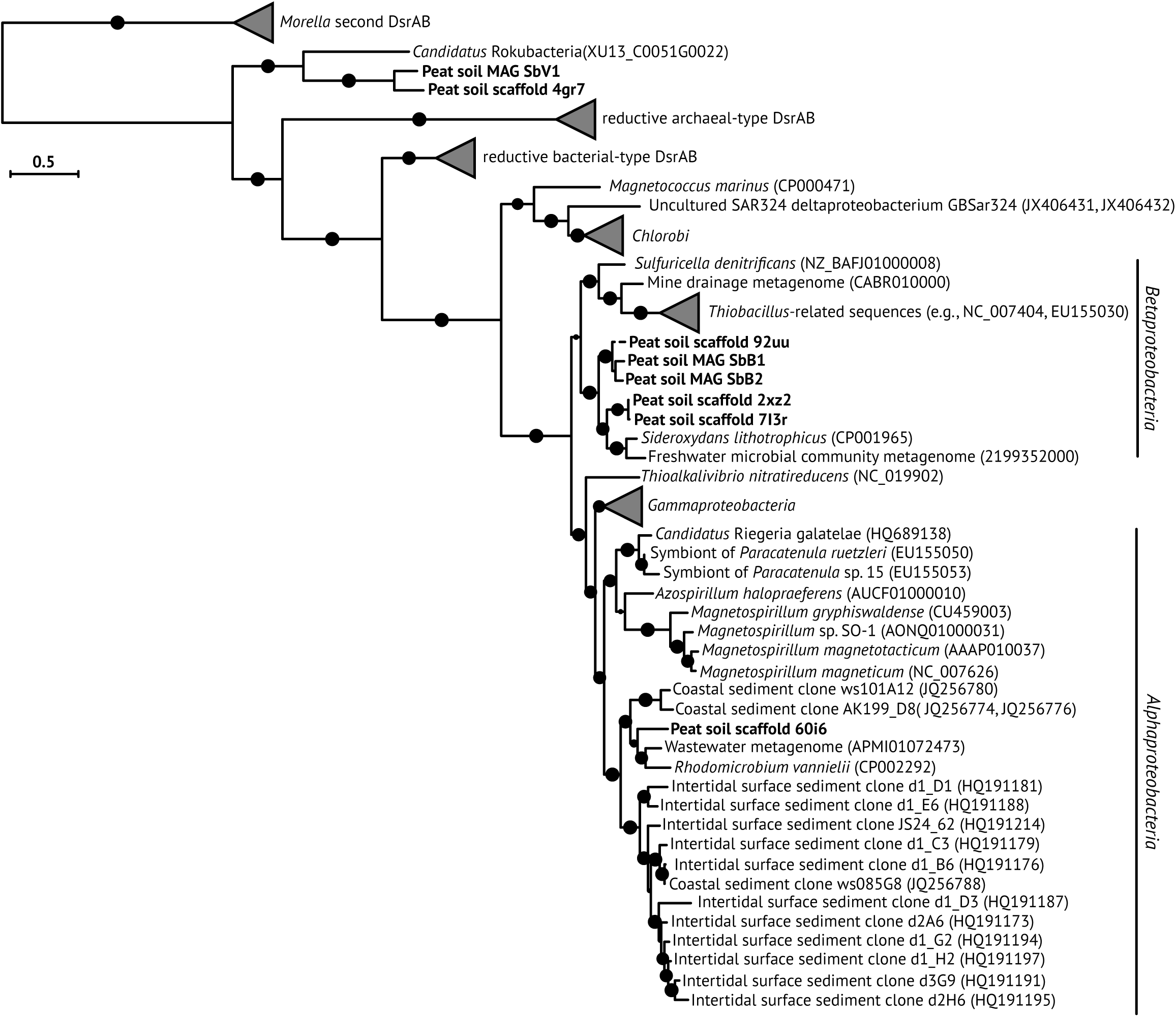
Oxidative bacterial-type DsrAB. Maximum likelihood phylogenetic tree was calculated by FastTree (LG model, 1000 resamplings) using a reference amino acid alignment with oxidative bacterial-type DsrAB indel positions removed (Müller *et al.*, 2015). Branch supports equal to or greater than 0.9 are indicated by black circles. DsrAB sequences from MAGs and scaffolds are marked in bold. Dashed branches represent partial DsrAB sequences on scaffolds.

**Supplementary Figure S3.**
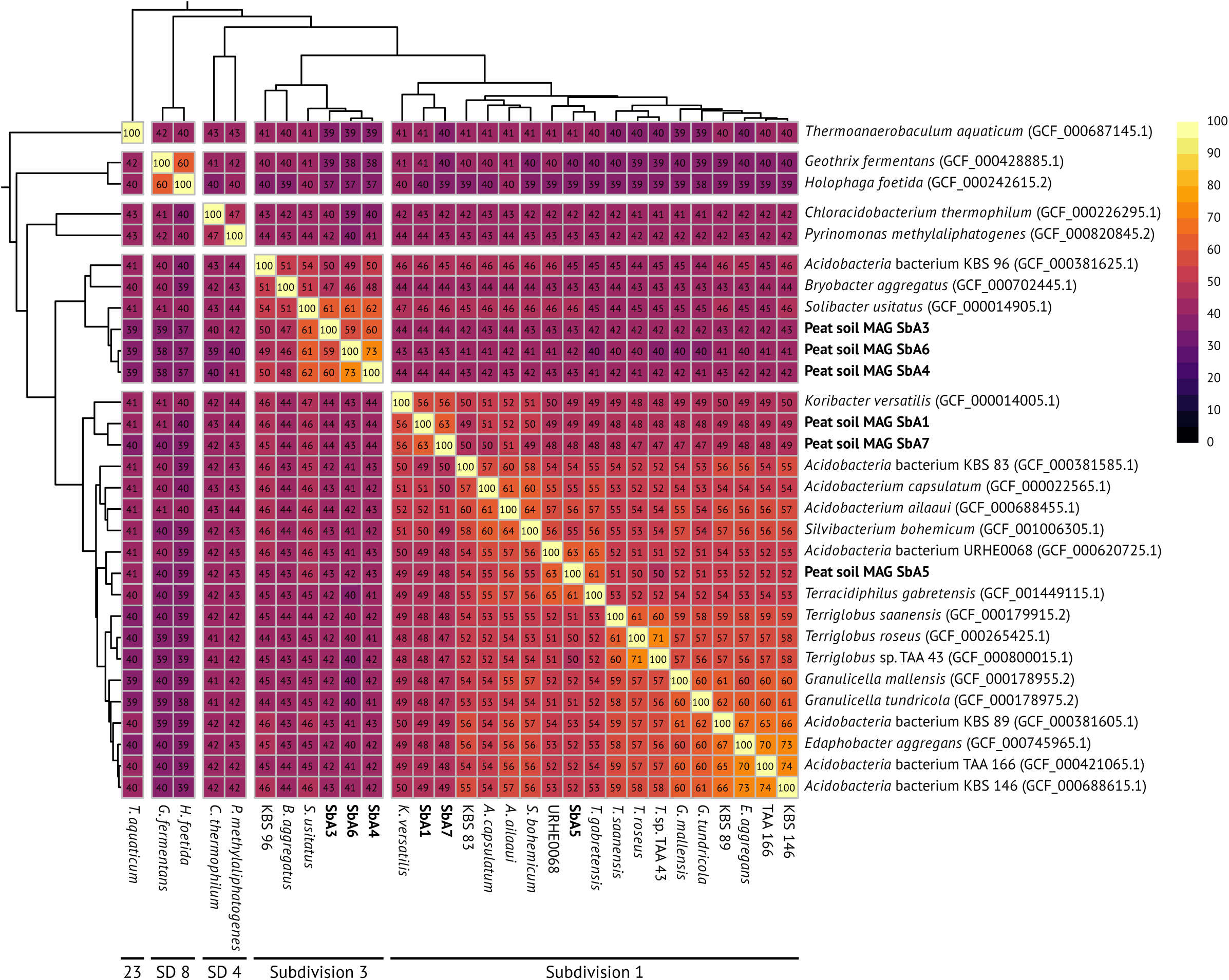
Phylogenomic tree and pairwise average amino acid identities of *Acidobacteria* genomes and MAGs. NCBI assembly accessions are given in parentheses. Novel sequences from this study are marked in bold. *Acidobacterial* subdivisions are given below. Dendrograms are a phylogenomic tree calculated with phylobayes from a checkM-produced and-filtered amino acid alignment. Allbranches are supported >0.9. Genomes assemblies from the *Firmicutes, Proteobacteria*, and *Verrucomicrobia* were used as outgroup. MAG SbA2 has an AAI of 37 – 49% to the other genomes and MAGs but was not included in the figure because it lacks the marker genes used for the phylogenomic tree.

**Supplementary Figure S4.**
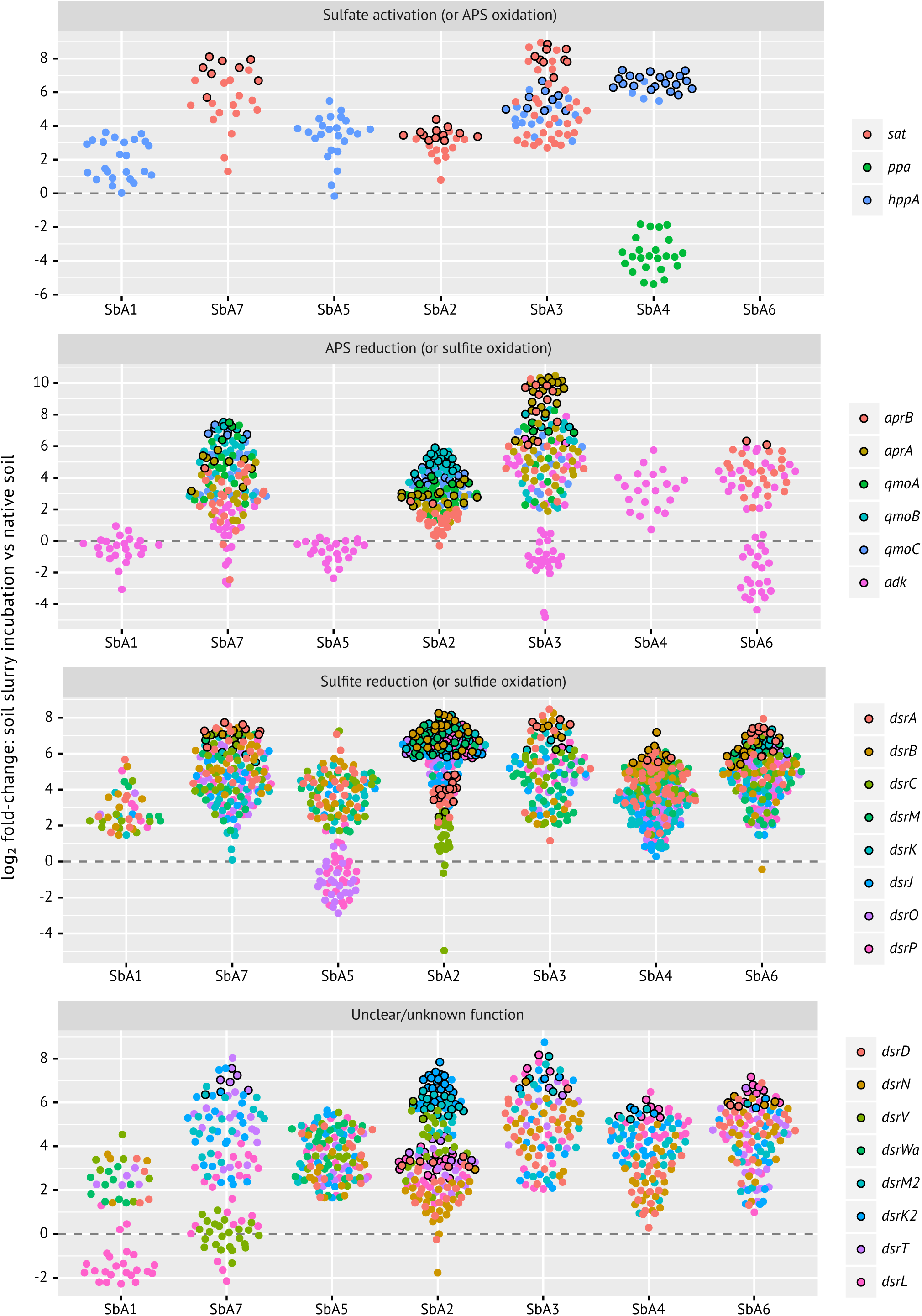
Beeswarm plots of expression change of dissimilatory sulfur metabolism genes in anoxic peat soil microcosms. Fold-changes are calculated by pairwise comparisons between replicate metatranscriptomes of the native soil and of each incubation regime and time point. Significant (*p*<0.05) changes are highlighted by back circles.

